# Dark state-mediated photobleaching in mCherry-based red fluorescent proteins

**DOI:** 10.64898/2026.01.21.700914

**Authors:** Premashis Manna, Mark A. Hix, Srijit Mukherjee, Alice R. Walker, Ralph Jimenez

## Abstract

Developing bright and photostable red fluorescent proteins (RFPs) is one of the ‘holy grails’ of the protein engineering community. Despite several attempts, finding such fluorescent proteins (FPs) has remained elusive. One bottleneck to engineering next generation RFPs is our lack of understanding of non-fluorescent or dark state properties in such constructs. Here, we develop a theoretical and experimental framework that describes how photobleaching decays in FPs relates to dark state conversion and ground state recovery. Our systematic photophysical investigation of mCherry and mCherry-d, an RFP with enhanced dark state behavior, showed the presence of photodestructive dark states in such FPs. Molecular dynamics simulations reveal enhanced fluctuation around the imidazolinone-end of the chromophore in mCherry-d, potentially facilitating conversion to non-fluorescent states. Collectively, this work quantifies dark state kinetics and gives insights into engineering dark states in RFPs to develop bright yet photostable molecular probes.

---

Fluorescent proteins (FPs) have gained widespread use as molecular probes in fluorescence microscopy. In the last few decades, FPs have been genetically engineered to enhance traits such as brightness, photostability, fluorescence color, and maturation time.^1^ This iterative process has yielded a diverse range of improved FPs for various applications.^2–6^ The photophysical properties of such FPs are sometimes as good as those of many small molecule dyes. For instance, the brightness of mTurquoise2,^6^ mNeonGreen,^3^ and mScarlet3,^7^ those emit in the blue, green and red spectral regions, respectively, have similar brightness of many small molecule dyes like Alexa, Atto or Janelia Fluor dyes.^8^ However, the photostability of the FPs, particularly for the red fluorescence proteins (RFPs) is still far from optimal.^9,10^ For instance, tetramethylrhodamine, one of the most used small-molecule dyes in laser spectroscopy has a *ϕ*_*PB*_ value of 3.3x10^-7^. On the contrary, the *ϕ*_*PB*_value for super photostable FPs like StayGold found in the FPbase^11^ is three orders of magnitude higher (*ϕ*_*PB*_ ∼ 10^-4^) under comparable illumination.^9,12^ This indicates that there is a large room for improvement in the photostability of FPs.

It has been reported that the fluorescence brightness and photostability of FPs are inversely-correlated.^13–15^ However, recent reports of bright and photostable green and yellow FPs challenge this trade-off in FP engineering.^4,12,16^ For example, StayGold, derived from the jellyfish *Cytaeis uchidae*, and its monomeric counterparts display ∼20-fold higher photostability than EGFP, yet maintain high molecular and cellular brightness.^12,16,17^ By employing a high-throughput single-cell screening platform, Lee et al. reported bright and photostable mGold2t and mGold2s which are ∼25-fold higher photostable than popular yellow fluorescent proteins (YFPs) such as mVenus and mCitrine.^4,18^ These improvements in photostability were not achieved at the cost of reduced brightness. However, a similar breakthrough is yet to occur in RFPs. Multiple attempts to engineer bright and photostable FPs have resulted in deterioration of one of the properties.^5,7,19,20^ For instance, although recent publications of mScarlet3-S2 or mScarlet3-H report higher photostability, these FPs sacrifice the brightness significantly relative to their predecessors.^21,22^

One of the main bottlenecks to improving the photostability of the RFPs while maintaining their brightness is our dearth of understanding possible photobleaching mechanisms and how they are controlled by photo-induced non-fluorescent or dark states.^23^ Previous studies indicate that these dark states might both be photoprotective and photodestructive.^23,24^ Population transfer from the bright state to a dark state is known as dark state conversion (DSC), whereas the relaxation from dark state to ground state is termed as ground state recovery (GSR) process (Figure 1a).^23,25,26^ Ensemble fluorescence measurements of closely related RFPs by our lab revealed the complex kinetics of these processes over six orders of magnitude in time.^23^ Photobleaching kinetics of TagRFP, mKate, and several FPs from the mFruit series^27^ were fit to a tri-exponential function. The fast component of the decay ranging from *μs* to ms was attributed to DSC. On the other hand, the slower weighted bi-exponential component (ms-s) was assigned to photodestruction. However, the explicit connection between microscopic parameters such DSC, GSR and photobleaching rates to the macroscopic observable such as photobleaching decays, was not made.

**Figure 1.**
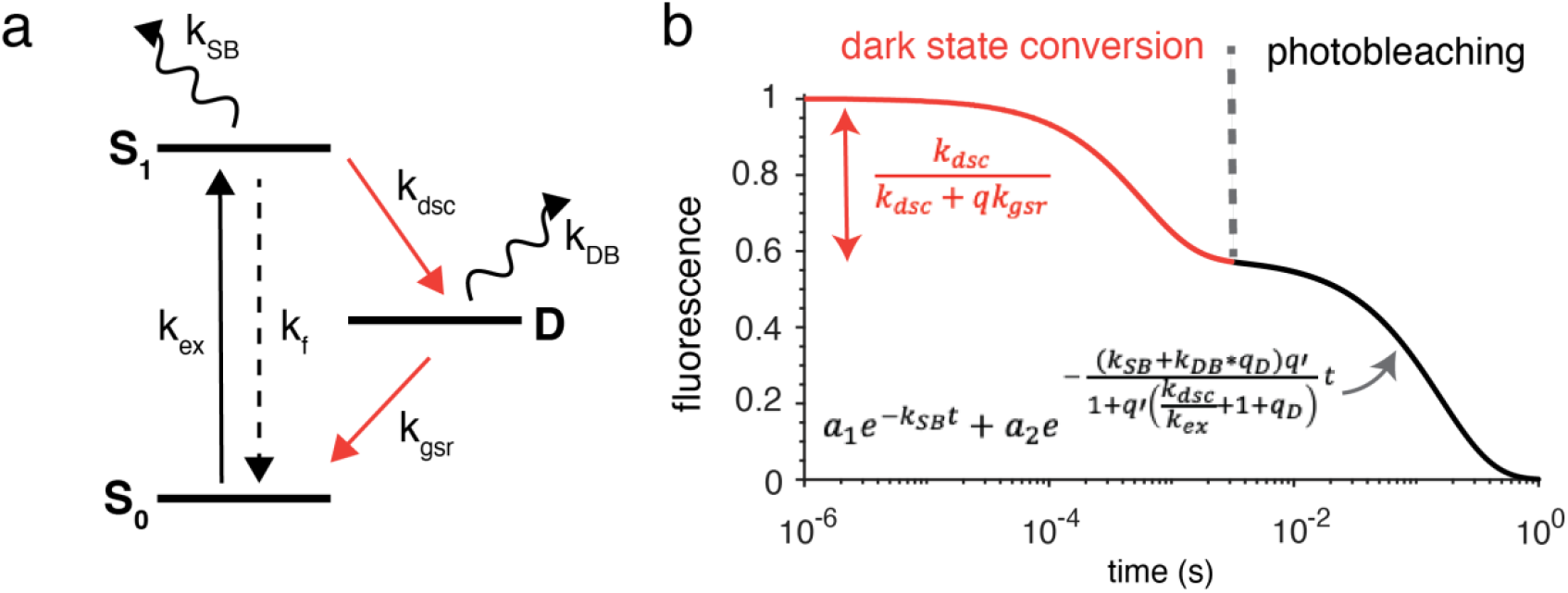
Kinetic model for dark state conversion and photobleaching. (a) A three-state photophysical model of the fluorescent proteins with the ground (S_0_), excited (S_1_), dark states (D), along with the relevant photokinetic parameters: the excitation rate (*k*_*ex*_), fluorescence rate (*k*_*fl*_), dark-state conversion rate constant (*k*_*dsc*_), ground-state recovery rate constant (*k*_*gsr*_), and the photobleaching rate constants from the bright and dark states (*k*_*SB*_ and *k*_*DB*_). (b) Typical fluorescence trace obtained from the model as described in (a), where the fast decay within milliseconds (orange line) is associated with the reversible dark state conversion and the slower decay (black line) is due to irreversible photobleaching. The relevant equations used to extract the *k*_*dsc*_ and photobleaching decays (*k*_*SB*_ and *k*_*DB*_) are also shown in the plot. Here 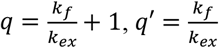 and 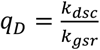.

In this work, first, we derive an analytical expression that describes how the time-constant and amplitude of the sub-ms fluorescence decay in FPs are governed by DSC, GSR, and excitation rates. Later, we adopt the photokinetic modeling, previously used in small-molecule chromophores to reveal the interdependence of these parameters on bi-exponential photobleaching observed in FPs.^28–30^ Second, we employ this method to extract kinetic parameters of the mCherry^27^ and mCherry-d, an RFP that we selected using a lifetime and photostability-gated microfluidic cell sorting system developed by Dean et al.^15^ mCherry and mCherry-d are closely related and have similar molecular brightness (*i*.*e*., extinction coefficient × fluorescence quantum yield) yet display contrasting dark state and photobleaching properties upon illumination in kW/cm^2^ regimes. Our analysis indicates a photodestructive dark state in mCherry-d. Intensity-dependent photobleaching measurements of the RFPs reveal that higher illumination intensity amplifies both kinetics and amplitude of dark-state mediated photobleaching. Finally, to investigate the molecular origin of such dark states, we performed all atom explicit solvent molecular dynamics (MD) simulations. We discover a significant destabilization of the phenolate ring (P-ring) and imidazolinone ring (I-ring) of mCherry-d chromophore relative to its precursor, mCherry. This is particularly noteworthy for the I-ring of the chromophore, as revealed by a large deviation of the I-ring dihedral angle from zero as well as enhanced fluctuations. We present analytical formulae for quantifying dark state properties from time-resolved fluorescence measurements, which are pivotal for designing high-throughput selection strategies for improved photophysical parameters. Identification of the molecular origin of dark states will guide the design of FP libraries either to stabilize or eliminate these states for the application of localization-based super-resolution microscopies or widefield/confocal imaging, respectively.^31^

## Modeling the dark state dynamics and photobleaching

We describe the photokinetics of the RFPs by a three-state photophysical model as described in Figure 1a. Upon illumination, the chromophore of the RFP is excited from the ground state (S_0_) to the excited state (S_1_). From S_1_, the molecule can be trapped in a dark state (D) and subsequently can relax back to S_0_. The rate constants for entering and exiting from D are *k*_*dsc*_ and *k*_*gsr*_, respectively. The model assumes that photobleaching occurs either from the singlet excited state or dark state with characteristic rate constants of *k*_*SB*_ and *k*_*DB*_, respectively. Here, the time-constant of any process is taken as the reciprocal of the relevant rate constant (i.e., *τ*_*dsc*_ = 1/*k*_*dsc*_).

The typical rate constants for excitation (*k*_*ex*_) and fluorescence (*k*_*f*_) are on the order of a few MHz (for irradiance regimes nearing optical saturation, i.e. ∼kW/cm^2^ intensity) and 100s of MHz (∼ns), respectively (see SI Sec S1). On the other hand, the photobleaching, DSC, and GSR processes are significantly slower - in the range of a few kHz (∼ ms) to tens of kHz (50 μs).^23,25,26^ These slower kinetics are consistent with the assumption that the dark and bleached states are effectively non-absorptive to the excitation photons driving the S_0_–S_1_ transition. Therefore, the kinetics of S_0_ and S_1_ states can be separated from the dark state conversion or photobleaching processes. In this limit, a rapid equilibrium is assumed between S_0_ and S_1_ states, i.e., *k*_*f*_[*S*_1_]_0_ = *k*_*ex*_[*S*_0_]_0_. Here, [*S*_0_]_0_ and [*S*_1_]_0_ are the equilibrium populations of the corresponding states. The validity of this approximation is justified by numerically solving for the exact populations in ground and excited states (SI, Sec S1).

At the illumination intensities employed here (∼kW cm^−2^), the characteristic dark-state conversion (DSC) time constant (tens of microseconds) can be an order of magnitude shorter than that of irreversible photobleaching, which typically occurs on timescales of tens of milliseconds. Consequently, over short observation windows (<1 ms), photobleaching can be treated as negligible. This approximation led us to derive an analytical expression of the excited state population as,

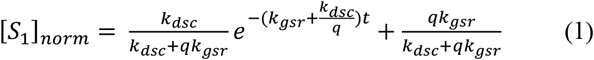

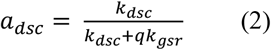

where, 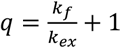 and *a*_*dsc*_ is the DSC amplitude. Detailed derivation of these equations and their validation from numerical simulation is given in SI, Sec S2.

Figure 1b shows a representative fluorescence bleaching trace simulated using the three-state model described in Figure 1a. The corresponding rate equations were solved numerically, and the normalized excited-state population (S_1_) was used as a proxy for the fluorescence signal. When plotted on a logarithmic time axis, the fluorescence trace exhibits an initial rapid decay on the sub-millisecond to ∼1 ms timescale, followed by a quasi-steady plateau and a subsequent slow exponential decay spanning the millisecond-to-second regime. The early-time decay arises predominantly from dark-state conversion (orange trace in Figure 1b), with negligible contribution from irreversible photobleaching over this interval. In contrast, the long-time decay beyond ∼1 ms is dominated by irreversible photobleaching processes (black trace in Figure 1b).

As indicated by Eqn. 1, at short timescales (within 1 ms) where photobleaching is negligible, the fluorescence decay is well described by a single-exponential function. Given that either *τ*_*gsr*_ or *τ*_*dsc*_ is known from independent time-domain experiments,^25^ the remaining parameter can be extracted from fits to the experimental fluorescence decay (Fig 2). In this work, *τ*_*gsr*_ for the RFPs was determined from independent time-domain measurements (Fig 2a-b) and subsequently used to extract *τ*_*dsc*_. We termed the amplitude of this fast decay as “DSC amplitude” (*a*_*dsc*_, Eqn. 2). In SI Sec S3, we show how *a*_*dsc*_ depends on various other kinetic parameters of our three-state model. In theory, both DSC amplitude 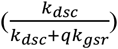 and the decay exponent 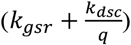 can be used to determine DSC time constants. In practice, however, the decay exponent is highly sensitive to a small number of early-time data points and is therefore prone to sampling errors. By contrast, *a*_*dsc*_ can be extracted more robustly, providing a reliable measure of DSC time constants over a broader parameter range, as demonstrated in SI Section S3 (Figure S3.1)

**Figure 2:**
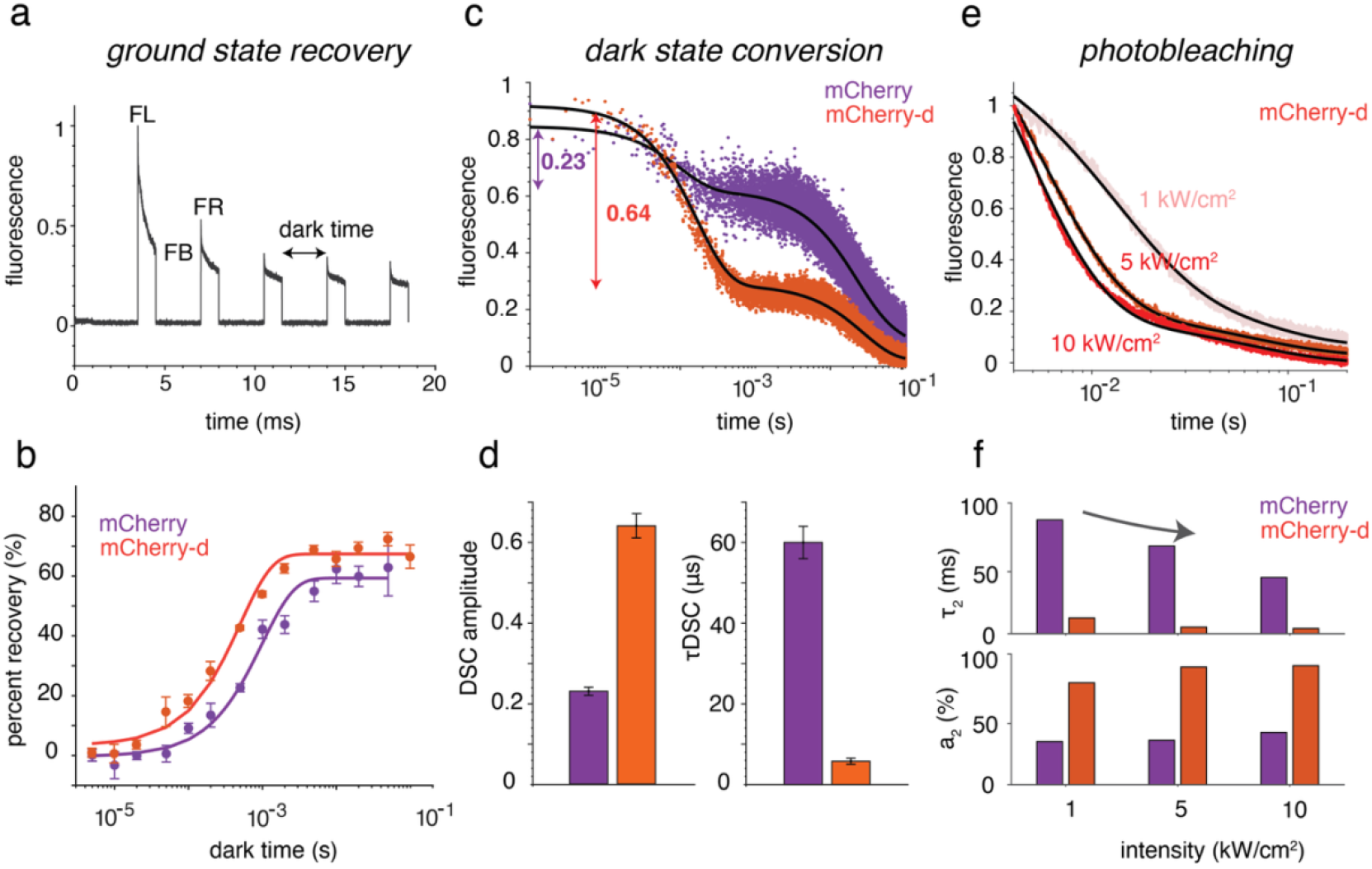
Dark state and photobleaching properties of mCherry and mCherry-d. (a) Typical fluorescence signal obtained from a FP sample in a ground state recovery (GSR) experiment. Cells expressing RFPs are excited with a 2 ms pulse and varying inter-pulse delays or dark times. Values of FL, FB, and FR are extracted from these experiments to calculate percent recovery employing Eqn. 4. (b) The percent recovery vs. dark times (dots) is fit to a single exponential (solid lines) to obtain GSR time constants. The error bars are standard deviations from three technical replicates. (c) Fluorescence traces demonstrating higher amplitude dark state conversion in mCherry-d (orange) compared to mCherry (purple) at 5 kW/cm^2^. While the fraction of fluorescence drops within 1 ms for mCherry is only 0.23, it is 0.64 for mCherry-d. The black lines are exponential fits. (d) DSC amplitude and time constants (*τ*_*DSC*_) for mCherry and mCherry-d are shown in purple and orange bars, respectively. The error bars are standard deviations from at least four independent measurements. (e) Normalized photobleaching decays of mCherry-d at 1, 5, and 10 kW/cm^2^. For each sample, the decays at different intensities are globally fit with a bi-exponential function (black lines). In the fit, *τ*_1_ is kept as a shared variable, whereas the amplitudes (*a*_1_, *a*_2_) and *τ*_2_ are intensity data specific. (f) Amplitudes and time-constants of the second component of mCherry (purple) and mCherry-d (orange) at different intensities.

It is noteworthy that although Eqn. 1 can be used to estimate both *τ*_*gsr*_ or *τ*_*dsc*_ from experimental fluorescence traces by fitting, such analyses generally incur substantial uncertainty. Consequently, independent measurements are required to reliably quantify these rates. For instance, in this previous report,^25^ we obtain GSR rates from time-domain measurements and then utilize the GSR rates to extract *τ*_*dsc*_ in a phase-based frequency-domain measurement. In a separate report, *τ*_*gsr*_ were determined from off-times of single molecule blinking in RFPs using TIRF-based measurements, while *τ*_*dsc*_ was extracted by solving the corresponding eigenvalue equations and fitting the resulting model to the ensemble fluorescence decay in bacteria.^26^ In both cases, however, an explicit dependence of dark-state kinetics on the fluorescence decay was not considered. This omission is justified, for example in the case the irradiation levels in single-molecule experiments (∼W cm^−2^) are approximately three orders of magnitude lower than those used here, resulting in excitation rates that are not rate-limiting; under these conditions, the dominant timescale reflects depopulation of the dark states, manifested primarily as off-times in single-molecule trajectories corresponding to *τ*_*gsr*_ . In contrast, the present work explicitly reveals and quantifies the dependence of fluorescence decay on dark-state kinetics under high-irradiance conditions.

Now, we adopt an analytical expression for photobleaching decays as solved for small-molecule dyes with a similar three-state (Figure 1a).^30^ In this case, by applying the rapid equilibrium approach as discussed above, the fluorescence decay (*F*(*t*)) can be expressed as,

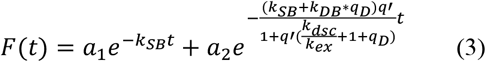

where *a*_1_ and *a*_2_ are constants, 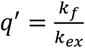 and 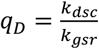. The details of the derivation are presented in SI, Sec S4. As evident from Eqn. 3, the photobleaching decays are biexponential, in agreement with our experimental results. The first term is due to the decays from solely S_1_ state (i.e., *k*_*SB*_), while the second term involves photobleaching rate constants from both the S_1_ and D states (i.e., *k*_*SB*_ and *k*_*DB*_). The second component also depends on the excitation rate (*k*_*ex*_), DSC rate constants (*k*_*dsc*_), and ratio of DSC and GSR rate constants (*q*_*D*_).

## Photophysical investigations of mCherry variants

Next, we apply the theoretical formulation developed above to experimental data to extract relevant kinetic parameters. For this, we performed systematic photophysical characterization of mCherry and mCherry-d (“d” for dark), a mutant of mCherry characterized by a higher amplitude of dark state formation. The mCherry-d mutant was developed via directed evolution of mCherry using a high-throughput microfluidic sorter as described in the SI (SI Sec S5) and here.^32^ Relative to mCherry, this variant contains three internal mutations (I161M, Q163M, and I197R) near the phenolate end of the chromophore. Also, it contains the K70R internal mutation near the imidazolinone end of the chromophore. These amino acid positions are reported to play important roles in brightness, maturation, and photostability in FPs.^2,27,33^ The full set of mutations in mCherry-d is provided in the SI (SI Table S1).

The photophysical parameters for mCherry and mCherry-d are displayed in Table 1. The blue shift of the absorbance maximum is consistent with the incorporation of the positively charged I197R mutation, as reported previously.^34^ MD simulations suggest the incorporation of I197R in mCherry-XL leads to the formation of multiple H-bonds which enhances the overall rigidity of the chromophore.^20,34^ The reduction of chromophore flexibility is consistent with the longer excited state lifetime in mCherry-d compared to its precursor. Although the excited state lifetime of mCherry-d is ∼30% longer than that of mCherry, they have very similar fluorescence quantum yields. Detailed photophysical characterizations, including the measurements of excitation/emission spectra, excited state lifetime, quantum yield and extinction coefficients are given in SI Sec S6.

**Table 1:** Photophysical properties of mCherry and mCherry-d variants.

| RFP | $\lambda_{abs}(nm)$ | $\lambda_{ems}(nm)$ | $\tau(ns)$ | $\epsilon(M^{-1}cm^{-1})$ | $\phi$ | $\tau_{dsc}(\mu s)$ | $\tau_{gsr}(ms)$ |
| --- | --- | --- | --- | --- | --- | --- | --- |
| mCherry | 587 | 610 | 1.72 | 74 | 0.25 | 60 | 1.0 |
| mCherry-d | 572 | 608 | 2.22 | 55 | 0.27 | 5.8 | 0.5 |

Under the experimental scheme presented in Figure 2a-b, yeast cells expressing mCherry and mCherry-d are excited with a pulse of 2 ms exposure time and varying inter-pulse delays (dark time) ranging from 5 *μs* to 100 ms in a custom-built inverted microscope.^25^ Figure 2a shows a typical fluorescence trace obtained from such excitation. Fluorescence intensities at points FL, FB, and FR in Figure 2a are extracted from such experiments to calculate the percent recovery (*PR*) employing,

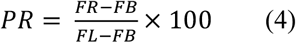

The PR values obtained from Eq. (4) were plotted as a function of the dark time and fitted to a single-exponential function to extract the GSR time constants. Using this analysis, we found that mCherry-d exhibits approximately two-fold faster recovery kinetics compared to mCherry, with time constants of 0.5 ms and 1 ms, respectively, suggesting that the depopulation from dark states into ground state is faster in mCherry-d. Additional details of the GSR measurements and analysis are provided in SI Sec S7.

We next focus on the dark state conversion (DSC) kinetics. Figure 2c-d demonstrates the starkly different dark state conversion in mCherry and mCherry-d. To extract DSC amplitude and time-constants, the normalized fluorescence decays from yeast cells are fit to a tri-exponential function. Although the DSC decay can be fit to a single-exponential function at short times (ref Eqn. 1), the presence of photobleaching components in observed fluorescence requires a tri-exponential fit. However, the faster *μs* component and the corresponding amplitude from these tri-exponential fits are taken as the DSC exponent and DSC amplitude. As discussed above, DSC amplitude is a better quantity to extract *τ*_*dsc*_ compared to the exponent. Therefore, DSC rate constants are obtained using Eqn. 2. *τ*_*gsr*_ and *q* are measured independently. The details of these measurements and analysis are given in SI Sec S8. We measure a DSC time-constant of (*τ*_*DSC*_, 60 μs) for mCherry, ∼10 fold-slower compared to mCherry-d (*τ*_*DSC*_, 6 μs). This difference is evident from the pronounced fluorescence drop within the first millisecond of illumination for mCherry-d compared to mCherry (Fig. 2c). Moreover, the DSC amplitude increases from 0.23 in mCherry to 0.64 in mCherry-d, indicating a higher propensity of mCherry-d to populate dark states.^23^

Figure 2e shows the normalized photobleaching decay of mCherry-d expressed in yeast cells at 1, 5, and 10 kW/cm^2^ illumination intensities. For each sample and intensity, three independent measurements were taken. The initial sub-ms fluorescence decay is largely due to dark state conversion and therefore negligible for irreversible photobleaching analyses and fits. For each sample, the remaining background-corrected decays at different intensities are globally fit with a bi-exponential function of the form:

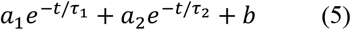

where, *a*_1_ and *a*_2_ are the amplitudes of the decay; *τ*_1_ and *τ*_2_ are photobleaching time-constants and *b* is a constant. As shown in Eqn. 3, the photobleaching decay can be modeled as bi-exponential where the first term depends only on photobleaching from S_1_ state (i.e., *k*_*SB*_), while the second exponent depends on *k*_*ex*_ which is related to laser intensity. Therefore, for each sample, decays with different intensities are globally fit with a bi-exponential function where *τ*_1_ is kept as a shared variable. On the other hand, the amplitudes (*a*_1_, *a*_2_) and *τ*_2_ are kept as intensity data dependent. The fits are shown as black lines in Figure 2e.

Table 2 displays the decay constants and amplitudes obtained from such global fits. As expected, within each sample, the weighted bleaching time-constants (*τ*_*avg*_) become faster with increasing illumination intensity. However, at the same intensity, mCherry-d bleaches ∼3-fold faster than mCherry. For instance, at 5 kW/cm^2^, *τ*_*avg*_ for mCherry is 29 ms, whereas it is 11 ms for mCherry-d. The first bleaching time-constant (*τ*_1_) which is essentially reciprocal of *k*_*SB*_ is slower in mCherry-d (10 ms vs. 63 ms). However, the higher amplitude and faster time-constants of the second component (*a*_2_ and *τ*_2_) largely contributes to the enhanced photobleaching in mCherry-d.

**Table 2:** Photobleaching time-constants of mCherry and mCherry-d variants.

| | intensity (kW/cm<sup>2</sup>) | $a_1$ | $\tau_1$ (ms) | $a_2$ | $\tau_2$ (ms) | $\tau_{avg}$ (ms) |
| --- | --- | --- | --- | --- | --- | --- |
| mCherry | 1 | 67 | 10 | 33 | 87 | 35 |
|  | 5 | 66 |  | 34 | 67 | 29 |
|  | 10 | 60 |  | 40 | 43 | 23 |
| mCherry-d | 1 | 22 | 63 | 78 | 12 | 23 |
|  | 5 | 10 |  | 90 | 5 | 11 |
|  | 10 | 9 |  | 91 | 4 | 9 |

Figure 2f plots *a*_2_ and *τ*_2_ at different intensities for mCherry and mCherry-d. For both FPs, we find that the amplitude of *a*_2_ increases and the corresponding decay constants (*τ*_2_) becomes faster with increasing laser intensity. More interestingly, at the same intensity, *a*_2_ for mCherry-d is significantly higher and *τ*_2_ is faster than mCherry. For instance, at 5 kW/cm^2^, *a*_2_ for mCherry and mCherry-d are 34% and 90%, respectively. On the other hand, *τ*_2_ is ∼13-fold faster for mCherry-d (67 ms vs. 5 ms), resulting in a markedly faster overall decay. As the second component involves DSC and GSR rates (Eqn. 3), these results indicate that the accelerated photobleaching decay in mCherry-d is mediated by dark state dynamics. Together, these kinetic analyses demonstrate that the enhanced accumulation of dark-state populations in mCherry-d renders it more susceptible to irreversible photobleaching originating from these states, which are therefore photodestructive. This is consistent with our previous report where we showed faster photobleaching in mCherry under a continuous-mode illumination compared to a pulsed-mode illumination condition, indicating a photodestructive dark states.^23^ On the contrary, several eqFP578-based RFPs such as TagRFP^23^ and FusionRed^24^ mutants are reported to have photoprotective dark states.

## Possible molecular origins of enhanced dark state behavior

The discovery of photodestructive dark states in mCherry mutants motivates us to investigate their possible molecular origin. Although such photophysics is inherently governed by excited state manifold of the FPs, conformational heterogeneity in the electronic ground-state revealed by molecular dynamics simulations provides valuable mechanistic insights. Therefore, we performed all-atom MD simulations with AMBER and explicit solvent (TIP3P) on mCherry and mCherry-d starting with the reported high-resolution crystal structure of mCherry (PDB, 2H5Q). Further simulation details are presented in SI Sec S10. The simulations suggest that the Q163M substitution in mCherry-d disrupts the hydrogen bonding (H-bonding) network at the phenolate end of the chromophore, while K70R and I197R mutations alter the H-bonding network around the imidazolinone end of the chromophore. For instance, in mCherry-d, the chromophore has increased hydrogen bonding with R70 (+36% vs 70K in mCherry), but decreased hydrogen bonding with M163 (-23% vs 163Q in mCherry). In addition, R197 in mCherry-d forms increased hydrogen bonds with the β-barrel backbone compared to I197 in mCherry, (e.g., a triad with K/R70 and E148) displacing and stabilizing the chromophore despite not directly hydrogen bonding with it. These residues are highlighted in Figure 3a, and the associated disruption of the H-bonding network is demonstrated in Figure S10. Collectively, these mutations reposition the chromophore toward R70 and promote altered dihedral twisting. Similar trends were observed in our previous systematic study that led to the development of mCherry-XL, the brightest mCherry mutant reported to date, where decreased chromophore and protein flexibility correlated with enhanced fluorescence brightness along a fluorescence-lifetime trajectory.

**Figure 3.**
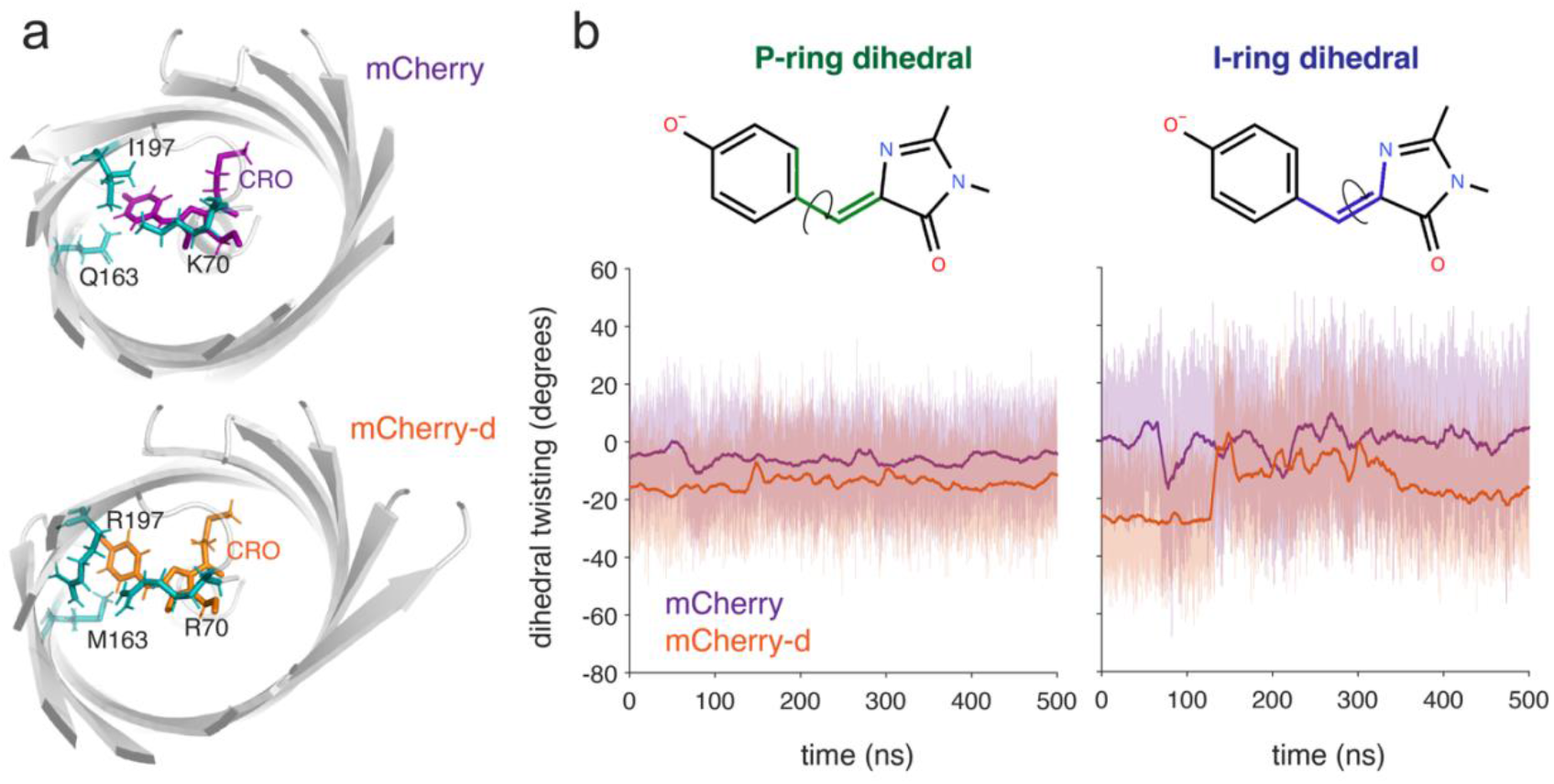
Molecular origin of dark states in mCherry-d. (a) Key residues responsible for the disruption of hydrogen bond networks in mCherry (up) and mCherry-d in a mCherry crystal structure (PDB 2H5Q). The mutations in mCherry-d are introduced in-silico. The chromophores are shown in purple for mCherry and orange for mCherry-d. The FP barrel structures are shown in grey. (b) Phenolate-ring (left) and imidazolinone-ring (right) dihedral twisting as a function of time for mCherry (purple) and mCherry-d (orange) as obtained from MD simulations. The solid lines are the moving average of the traces. The dihedral angles quantified are shown above for each plot.

Altered hydrogen bonding at the phenolate end also affects dihedral angles along the chromophore bridge. Increased rotational freedom facilitates access to nonradiative decay pathways in the excited state, which is reflected in enhanced ground-state twisting and reduced chromophore planarity^35,36^—features experimentally associated with lower fluorescence quantum yield.^20^ Figure 3b displays the P- and I-ring dihedral for mCherry (purple) and mCherry-d (orange) within 500 ns. Wild-type mCherry demonstrates I- and P-ring dihedral twisting from -20 to +20 degrees, centered at 0 degrees. mCherry-d, however, has P-ring rotations from -40 to +20 centered at -18 degrees. We correlate this to the Q163M mutation and loss of hydrogen bonding at the phenolate end of the chromophore. Previously, Regmi and co-workers used fixed-charge, explicit solvent all-atom MD simulations to show that the Q163M mutation alters the oxygen diffusion channels in mCherry.^37,38^ This resulted in a significantly higher rate and number of oxygen molecules entering the protein barrel compared to wild-type mCherry contributing to an enhanced photobleaching. Similar oxygen-dependent photobleaching may also operate in mCherry-d.

We also note an interesting large shift in the I-ring dihedral, which fluctuates from -40 to +40 degrees and is centered at ∼-20 degrees in mCherry-d. We attribute this to these mutations - K70R and I197R, which alter the chromophore cavity such that the alpha helix core and the chromophore cavity are destabilized. I-twisting is associated with nonradiative decay for HBDI-, but in an FP, the I-ring is bound on either side to the protein matrix.^39,40^ This suggests the possibility that the observed dark state is associated with a twisted I-ring and thus photodestructive, but potentially stable as it cannot complete isomerization due to chromophore-protein interactions. However, more work is needed to further characterize this possibility.

In conclusion, the work presented here provides a theoretical framework to extract dark state conversion rates from fluorescence measurements. We showed that both DSC and GSR contribute to the fast initial fluorescence drop in mCherry-based RFPs and the amplitude of DSC is a robust parameter to quantify *τ*_*dsc*_. This method could easily be extended to other FPs that emit at different wavelengths, such mGold2s, where a significant photoprotective dark state might be present.^4^ Employing analytical derivation and numerical simulations, we explain the bi-exponential nature of photobleaching decays in these FPs and how it relates to the dark state properties. By systematic photophysical investigations of mCherry and mCherry-d, we conclude the existence of dark-state mediated photobleaching in such RFPs. This behavior could also explain the higher magnetic field effects observed in mCherry-d fluorescence compared to mCherry, where dark states such as triplets can play a critical role.^41^ Intensity-dependent photobleaching measurements in these RFPs reveal that both the rate and amplitude of such decay are accelerated at high illumination intensity. Finally, with the help of molecular dynamics simulation, we discover a significant deviation of chromophore I-ring dihedral from zero and its enhanced fluctuation, which might contribute to the dark state behavior observed here. Collectively, the insights from this study will provide useful input for designing RFPs with better photostability and provide a foundation for engineering stable dark states, which is essential in many advanced localizations-based super-resolution microscopies such as MINFLUX.^42^

## Author Contributions

P.M., S.M. and R.J. conceptualized the study. M.A.H and A.R.W designed, performed and analyzed the simulations. P.M. designed and performed the experiments. P.M. derived the analytical equations and developed data analysis methods for this study. P.M., S.M., M.A.H, A.R.W., and R.J. wrote the manuscript.

## Notes

The authors declare no competing financial interest.

## Acknowledgement

P.M. was partially supported by his startup grant provided by The Ohio State University. S.M. was supported by the NIH/CU Molecular Biophysics Training Program (T32). This work was partially supported by the NSF Physics Frontier Center at JILA (PHY 2317149 to R.J.). R.J. is a member of the Quantum Physics Division of the National Institute of Standards and Technology (NIST). Certain commercial equipment, instruments, or materials are identified in this paper in order to specify the experimental procedure adequately. Such identification is not intended to imply recommendation or endorsement by the NIST, nor is it intended to imply that the materials or equipment identified are necessarily the best available for the purpose. We thank Prof. Amy E. Palmer (CU Boulder) and Prof. Sheng-Ting Hung (NSYSU) for valuable discussions. We also thank Prof. Steven G. Boxer and Prof. Soichi Wakatsuki (Stanford University), Dr. Irimpan Mathews (SLAC), and Nahal Bagheri (Stanford University) for insightful discussions.

## Supporting Information Available

Details of the photobleaching model, directed evolution of mCherry-d, measurement, and analysis of photobleaching, ground state recovery, dark state conversion and MD simulations can be found in the Supporting Information section of this article. This material is available free of charge via the Internet at http://pubs.acs.org.

