## Supplementary Info for "Dark state-mediated photobleaching in mCherry-based red fluorescent proteins"

**S1. Validity of rapid equilibrium approximation between the ground state and excited states.**

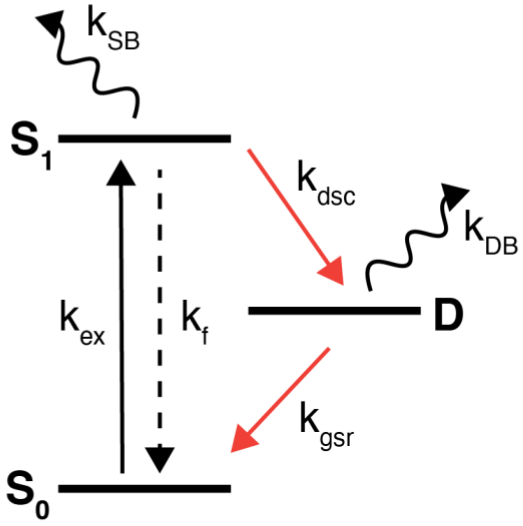


Figure S1.1: Kinetic model of the photophysical pathways.

Figure S1.1 shows the kinetic model of photophysical pathways. The proteins at the ground state are excited by a laser driving a population transition from $S_0$ state to $S_1$ state. The rate constant for excitation ($k_{ex}$) is given by:

$$k_{ex} = \frac{I_0 \sigma \lambda}{hc} \quad (1)$$

where, $I_0$, $\sigma$, and $\lambda$ are laser intensity, absorption cross section and laser wavelength. *h* and *c* are the Planck's constant and the velocity of light, respectively. $I_0$ is in the range of 1-10 kW/cm$^2$, $\lambda$ is 532 nm. Absorption cross-section ($\sigma$) is given as:

$$\sigma = 2.303 \times \frac{\epsilon_{532}}{N_A} \times 10^3 \quad (2)$$

where, $\epsilon_{532}$ is the molar extinction coefficient of the protein at 532 nm.

The molecule once excited to the $S_1$ state and relax back to the ground state in a radiative fashion or can get trapped in the long-lived dark state, D. The photobleaching can occur from the $S_1$ or D states with rate constants of $k_{SB}$ and $k_{DB}$, respectively. Here, we examine the nature of fluorescence trace in the case of zero photobleaching i.e. with $k_{SB} = 0, k_{DB} = 0$).

The population of different states can be obtained by solving the set of differential equations below:

$$\frac{d[S_0]}{dt} = -k_{ex}[S_0] + k_f[S_1] + k_{gsr}[D] \quad (3)$$

$$\frac{d[S_1]}{dt} = k_{ex}[S_0] - (k_f + k_{dsc} + k_{s1B})[S_1] \tag{4}$$

$$\frac{d[D]}{dt} = k_{dsc}[S_1] - k_{gsr}[D] \tag{5}$$

With laser intensity of 5 kW/cm$^2$, the excitation rate-constant for an RFP is typically in the range of few MHz. On the other hand, FPs fluoresces in the timescale of ns (~ 100 MHz). Compared to these fast timescales, the photobleaching and dark-state conversion rates are very slow, in the range of few kHz (~ ms) to tens of kHz (50 $\mu$s). Therefore, the kinetics of $S_0$ and $S_1$ state can be separated from the state D. Also, an equilibrium is established between $S_0$ and $S_1$. At this condition,

$$k_f[S_1]_0 = k_{ex}[S_0]_0 \tag{6}$$

$$[S_0]_0 = \left(\frac{k_f}{k_{ex}}\right)[S_1]_0 = q'[S_1]_0 \tag{7}$$

Where $q' = \frac{k_f}{k_{ex}}$ and $k_f = \frac{\phi}{\tau}$ with $\phi$ and $\tau$ are the fluorescence quantum yield and the fluorescence lifetime of the RFP, respectively. $[S_0]_0$ and $[S_1]_0$ are the equilibrium population of the corresponding states.

Again, at a steady state, there is no change in dark state population. Therefore,

$$\frac{d[D]}{dt} = k_{dsc}[S_1]_0 - k_{gsr}[D]_0 = 0 \tag{8}$$

$$[D]_0 = \left(\frac{k_{dsc}}{k_{gsr}}\right)[S_1]_0 = q_D[S_1]_0 \tag{9}$$

where, $q_D = k_{dsc}/k_{gsr}$ . Now, as there is no loss of population due to photobleaching, the total population must satisfy the condition below:

$$[S_0]_0 + [S_1]_0 + [D]_0 = 1 \tag{10}$$

Combining Eqn. 7, 9 and 10, we obtain,

$$q[S_1]_0 + [S_1]_0 + q_D[S_1]_0 = 1 \tag{11}$$

$$[S_1]_0 = \frac{1}{1+q+q_D} \tag{12}$$

Figure S1.2 shows the population of different states obtained from solving the set of differential equation above. The dotted lines are the steady state population obtained from Eqn. 7, 12, and 9. The population obtained from the steady state approximation agrees well with the results from numerical simulations and therefore validates our steady state approximation.

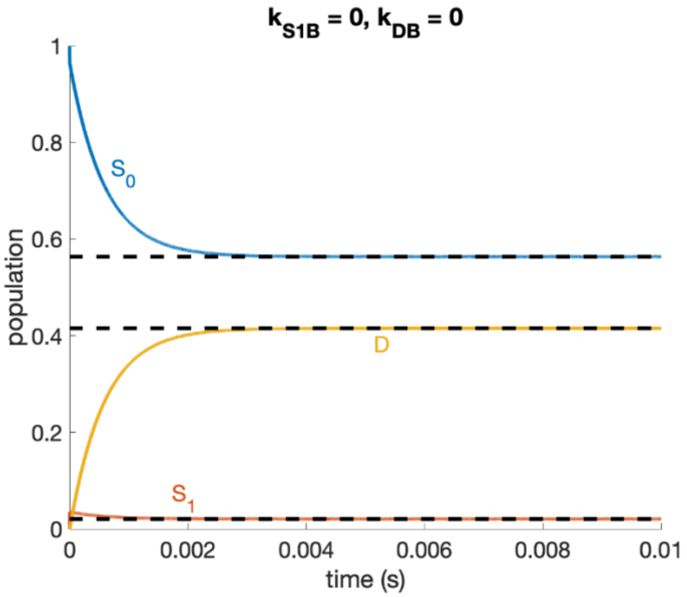


Figure S1.2: Population of different states obtained from the numerical solutions (solid lines) with zero photobleaching condition. The dotted lines are steady state population of $S_0$, $S_1$ and D obtained from Eqn. 7, 12 and 9, respectively.

**S2. Derivation of an analytical expression for DSC amplitude and exponent.**

For an analytical expression of DSC, we invoke the same rapid equilibrium approximation described and validated above.

Now, neglecting photobleaching at short sub-ms timescale and combining Eqn. 7 and 10, we have,

$$q'[S_1]_0 + [S_1]_0 + [D]_0 = 1$$
$$[D]_0 = 1 - [S_1]_0(1 + q') \quad (13)$$

Now, inserting Eqn. 13 in Eqn. 5 we obtain,

$$-(1 + q')\frac{d[S_1]_0}{dt} = k_{dsc}[S_1]_0 - k_{gsr}(1 - [S_1]_0(1 + q'))$$
$$-q\,\frac{d[S_1]_0}{dt} = k_{dsc}[S_1]_0 - k_{gsr}(1 - q[S_1]_0)$$
$$-q\,\frac{d[S_1]_0}{dt} = (k_{dsc} + qk_{gsr})[S_1]_0 - k_{gsr}$$
$$\frac{d[S_1]_0}{dt} = -\left(\frac{k_{dsc}}{q} + k_{gsr}\right)[S_1]_0 + \frac{k_{gsr}}{q} \quad (14)$$

Please note that, unlike the set of coupled differential equations in Eqn. 3-5, Eqn. 14 *only* describes the population of $[S_1]_0$, after it reaches a rapid equilibrium within sub-$\mu s$ timescale as described earlier. Therefore Eqn. 14 can be solved analytically with subject to the initial condition as follows.

As we are interested in the time-evolution of $[S_1]_0$ (which is proportional to the experimentally observed fluorescence), we define $t = 0$, after the population reaches the rapid equilibrium. At this short time (sub-$\mu s$), there is no sufficient build-up of the dark state population. Therefore,

$$[S_1]_0(t = 0) + [S_0]_0(t = 0) = 1 \quad (15)$$

Applying, Eqn. 7 in Eqn. 15, we have,

$$[S_1]_0(t = 0) = \frac{1}{q'+1} = \frac{1}{q} \quad (16)$$

By solving the Eqn. 14 with the initial condition shown in Eqn. 16, we have,

$$[S_1]_0 = \frac{k_{dsc}}{q(k_{dsc}+qk_{gsr})}\exp -(k_{gsr} + \frac{k_{dsc}}{q})t + \frac{k_{gsr}}{k_{dsc}+k_{gsr}q} \quad (17)$$

The normalized excited state population ($[S_1]_{norm}$), therefore, is given as,

$$[S_1]_{norm} = [S_1]_0 * q = \frac{k_{dsc}}{(k_{dsc}+qk_{gsr})}\exp -(k_{gsr} + \frac{k_{dsc}}{q})t + \frac{qk_{gsr}}{k_{dsc}+k_{gsr}q} \quad (18)$$

Figure S2 shows the comparison of exact numerical soln. obtained from Eqn. 3-5 and the approximate analytical soln. from Eqn. 18. It is evident that the results agree with each other very well validating the analytical approach.

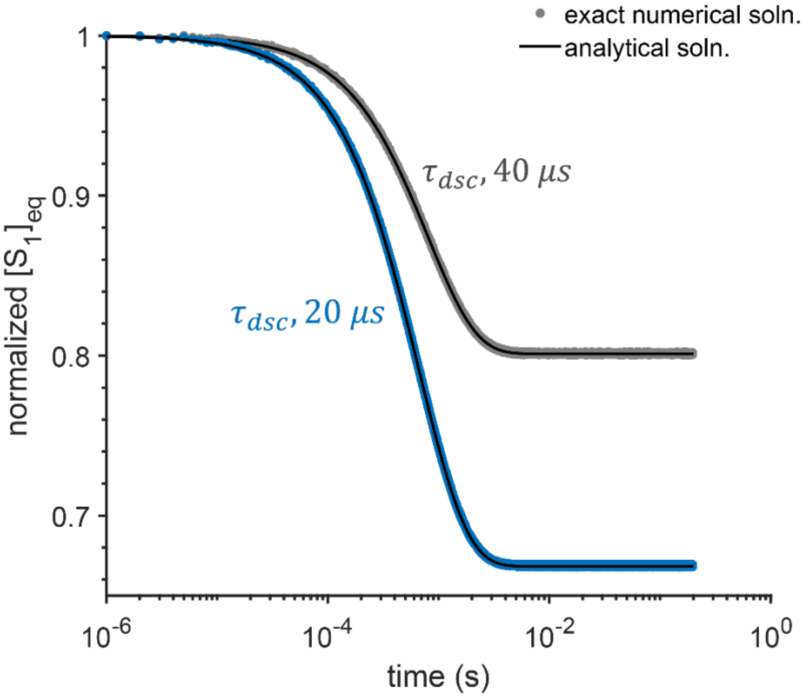


Figure S2: Comparison of exact numerical soln. obtained from Eqn. 3-5 and the analytical soln. from Eqn. 18.

Unlike the numerical solution, the analytical soln. clearly shows how the initial sub-ms decay of fluorescence is controlled by rates of DSC, GSR and the ratio of fluorescence emission ($k_f$) and laser excitation rates ($k_{ex}$). According to Eqn. 18, the rates of the fluorescence decay ($k_d$) in this short time-scale is

$$k_d = k_{gsr} + \frac{k_{dsc}}{q} \quad (19)$$

and amplitude of the fast component or 'DSC amplitude' ($a_{dsc}$) is,

$$a_{dsc} = \frac{k_{dsc}}{k_{dsc}+qk_{gsr}} \quad (20)$$

**S3. DSC amplitude ($a_{dsc}$) is a better quantity to extract $k_{dsc}$ than fluorescence rate constant ($k_d$).**

As both DSC amplitude (Eqn. 20) and fluorescence decay constant (Eqn. 19) depends on $k_{dsc}$, both can be utilized to extract DSC time constant ($\tau_{dsc}$). To examine which one is a better metric, we ran a series of numerical simulations and compare the results with the ground truth. We solve the set of Eqn. 3-5 numerically in MATLAB and fit the $[S_1]$ population (which is our proxy for fluorescence) obtained from simulation with a single-exponential function. In this model, we photobleaching was set to zero, and therefore, the $[S_1]$ population can be well fit a single-exponential function. The amplitude and exponent of the of the fit then utilized to extract $k_{dsc}$ using Eqn. 20 and 19, respectively.

The recovered $\tau_{dsc}$ which is the reciprocal of $k_{dsc}$ are plotted (blue dots) along the ground truth (black lines) in Figure S3. It shows that $\tau_{dsc}$ recovered from DSC amplitude (Figure S3.1 b) works well with a wide range of values while $\tau_{dsc}$ obtained from the rate agrees well with the ground truth only at very fast DSC rates (Figure S3.1 a). Therefore, we conclude that DSC amplitude is a better quantity to extract DSC time-constant than the fluorescence rate.

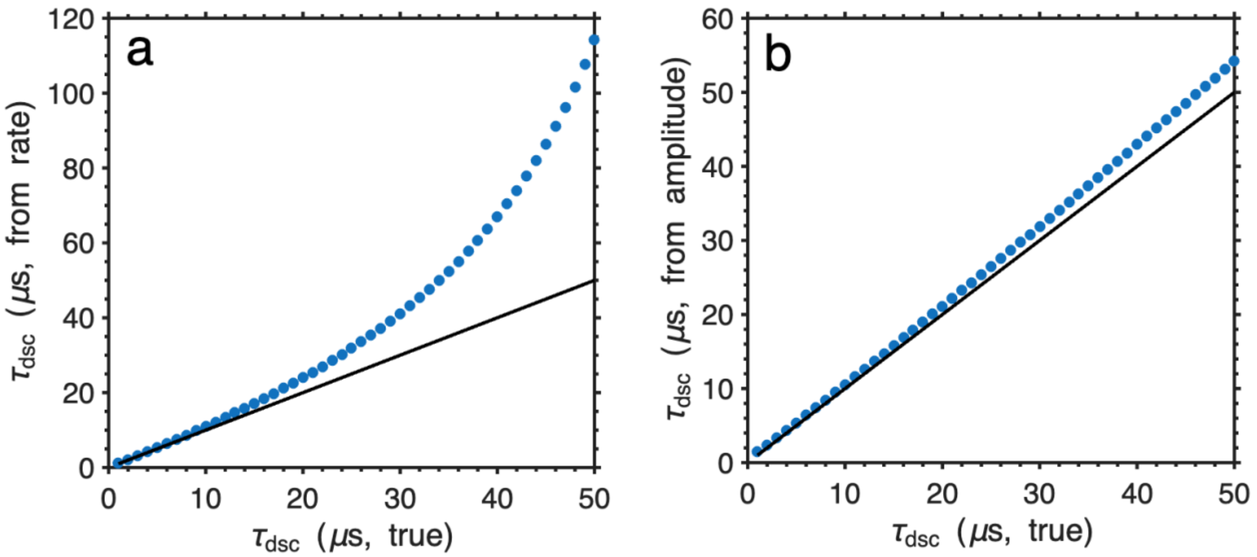


Figure S3.1: Comparison of $\boldsymbol{\tau_{dsc}}$ recovered from fluorescence decay exponent (a) and DSC amplitude (b) shown in blue dots. The ground truths are shown in black lines. The parameters used for the numerical simulations are, $\boldsymbol{k_{ex} = 1.47 \times 10^6\ s^{-1}}$, $\boldsymbol{k_f = 1.25 \times 10^8\ s^{-1}}$, $\boldsymbol{k_{gsr} = 2 \times 10^3\ s^{-1}}$, $\boldsymbol{k_{SB} = 0\ s^{-1}}$, $\boldsymbol{k_{DB} = 0\ s^{-1}}$.

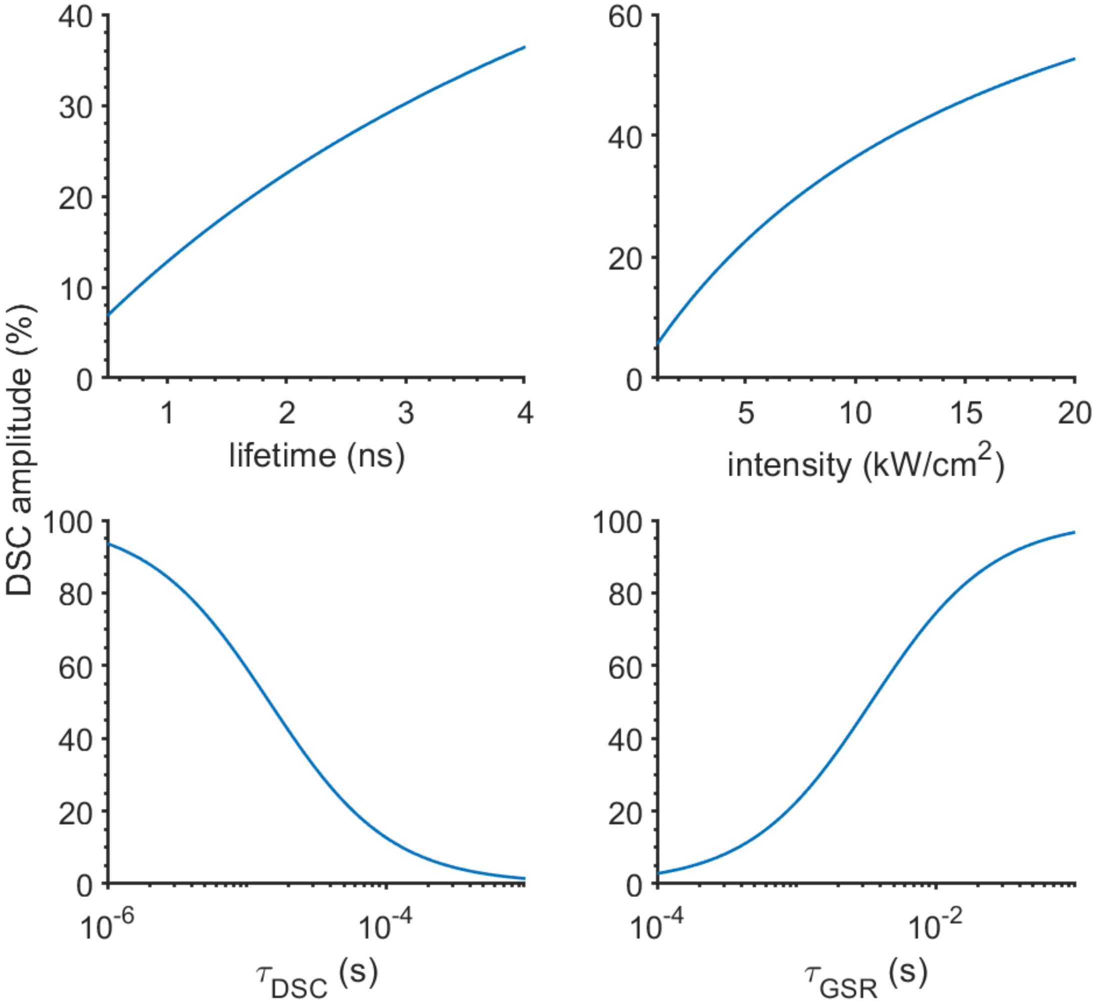


Figure S3.2: DSC amplitude as a function of different model parameters, such as excited state lifetime, laser intensity, $\boldsymbol{\tau_{dsc}}$ and $\boldsymbol{\tau\_gsr}$.

Figure S3.2 shows systematic investigation of different model parameters employing Eqn. 20. While one parameter varied, the others were kept constants. The plot shows that DSC amplitude increases as $\tau_{DSC}$ becomes faster. It also increases as the lifetime becomes longer and excitation intensity kept higher. On the other hand, DSC amplitude is smaller when ground state recovery is faster. Any condition that increases the population of dark states, the DSC amplitude becomes higher. In other words, this parameter is a good indicator of dark state population.

**S4. Theory of photobleaching decays from a three-state model.**

For the photobleaching decay kinetics, we adopt the results published by Wustner et al.[1] The authors showed how considering bleaching only from the singlet excited state (refer to Figure S1) leads to a single-exponential decay while a combination of photobleaching from $S_1$ and D states results in a bi-exponential decay. Here, we briefly discuss the analysis.

First, we examine the case where photobleaching occurs only from the excited state ($S_1$) and neglects any contribution from the dark state (D). This translates to a two-state model, and we employ the rapid equilibrium approach between $S_0$ and $S_1$ states. In this special case, the photobleaching is a first-order process and only takes place from $S_1$ with a rate constant of $k_{SB}$ and the fluorescence decay ($F_S(t)$) is given by,[1]

$$F_S(t) = m_1 \frac{q'}{1+q'} \exp\left(-k_{SB} \frac{q'}{1+q'} t\right) \quad (21)$$

where, $m_1$ is a constant and $q' = \frac{k_f}{k_{ex}}$. For a typical excitation rate and excited state lifetime of FPs, q >> 1, and therefore, the $\frac{q'}{1+q'}$ - term in the exponent and pre-exponent is close to unity. Therefore,

$$F_s(t) \approx m_1 * \exp\left(-k_{SB} * t\right) \quad (22)$$

Now, we consider the photobleaching both from the dark state and the excited state (i.e. we use the three-state model). In this case, by applying the rapid equilibrium approach as discussed above, the fluorescence decay ($F_D(t)$) can be expressed as,[1]

$$F_D(t) = m_2 \frac{q'}{1+q'(\frac{k_{dsc}}{k_{ex}}+1+q_D)} \exp\left(-\frac{(k_{SB}+k_{DB}*q_D)q'}{1+q'(\frac{k_{dsc}}{k_{ex}}+1+q_D)} t\right) \quad (23)$$

where, $m_2$ is a constant and $q_D = \frac{k_{dsc}}{k_{gsr}}$.

$m_1$ and $m_2$ in Eqn. 22 and 23 quantifies the fraction of the population photobleaches via. two state and three-state model systems, respectively.

In a case, where a fraction ($f$) of the population transferred to the dark state and the rest remain the bright state ($S_0$ and $S_1$ states). The total photobleaching, therefore, can be expressed as,

$$F(t) = (1-f)\exp(-k_{SB}t) + f \frac{q'}{1+q'(\frac{k_{dsc}}{k_{ex}}+1+q_D)} \exp\left(-\frac{(k_{SB}+k_{DB}*q_D)q'}{1+q'(\frac{k_{dsc}}{k_{ex}}+1+q_D)} t\right) \quad \text{or,}$$

$$F(t) = a\exp(-t/\tau_1) + b\exp(-t/\tau_2) \quad (24)$$

where, $\tau_1 = 1/k_{SB}$, $\tau_2 = \frac{1+q'(\frac{k_{dsc}}{k_{ex}}+1+q_D)}{(k_{SB}+k_{DB}*q_D)q'}$, $a$, $b$ are constants.

**S5. Development of mCherry-d mutant.**

The development of mCherry-d (or C4PB-12) from its precursor mCherry is shown in Figure S5.1 and also have been discussed in detail in here.[2]

Briefly, to enhance photostability in mCherry, we generated a fluorescent protein library – Kriek 1. The amino acid position targeted in Kriek 1 library were: V16, M66, W143, I161, Q163, I197, and A217. Position 143 and 163 was chosen to restrict entrance of oxygen inside the $\beta$-barrel. Although molecular oxygen ($O_2$) is essential for the auto-catalytic formation of chromophore in FPs,[3,4] it can also enhance the rate of irreversible photobleaching by reacting with the excited state chromophore. Chapagain et al. performed MD simulations on mCherry and Citrine (a yellow fluorescent protein) and suggested that the inferior photostability of mCherry might be due to its increased inter-strand dynamics between $\beta$-7 and $\beta$-10.[5] The rest of the positions were chosen as they are commonly mutated residues in the mFruit series of FPs.[6] Fluorescent colony screening of the library in E. coli showed that ~10% of the library was fluorescent. We sorted the library to enrich the population with higher photostability using our home-built microfluidic sorter.[7] This sorter-enriched population served as the template for our next round of targeted mutagenesis. The sorting protocol has been discussed in the next section.

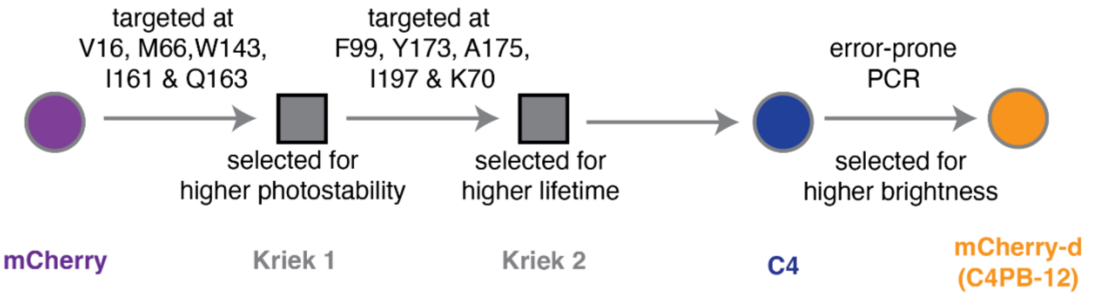


Figure S5.1: **Evolution trajectory of mCherry-d**. Directed evolution of mCherry-d from its precursor, mCherry. The mutants and fluorescent protein libraries are shown as filled circles and squares, respectively. The numbering of the amino acid residues is based on mCherry crystal structure (PDB: 2H5Q).

After the first round of targeted mutagenesis and subsequent enrichment of the photostability, we produced a population of lower photobleaching tendency. Unfortunately, the excited state lifetime of these mutants was significantly shorter than the parent mCherry, and therefore we suspected they would be of lower quantum yield. To improve the lifetime of those populations, we targeted the I197 residue, whose side-chain is reported to be capable of a $\pi$-stacking interaction with the chromophore.[8] Also, it was reported that mutations at position 70 are correlated with those at 197 (i.e. K70 "co-evolved" 197).[9] We therefore chose I197 and K70 as target residues for 2nd round of mutagenesis. Additional residues targeted in this phase of library generation were suggested by the MD simulations of mCherry by Regmi et al. [10] who explored the oxygen diffusion pathways into the $\beta$-barrel and identified F99, Y173, A175 as "gateway" residues. We hypothesized that rigidifying the barrel by reducing the motions of these dynamic residues would have a beneficial effect on the fluorescence quantum yield.

To isolate the mutants with higher lifetime, we grew our lifetime-enriched library on galactose-containing (for inducing expression in yeast) plates. Resulting yeast colonies were fluorescent and the lifetime for each colony was measured in a fashion similar to the in-flow phase fluorimetry, as described here.[11] From these colonies we identified the C4 mutant. The full set of mutations in C4 is: V16T, H17R, K70R, F99S, I161M, Q163M, A175W and I197R. It was found that mutations at position 16 were linked to a H17R (pointing outwards of $\beta$-barrel) mutation. This was not an intentional part of the Kriek 2 library and was due to an error in the primer design. The C4 mutant was promising as it showed improved excited state lifetime and photostability in the microfluidic screening. However, this mutant was dimmer than the parent mCherry. This dimness may be due to its lower extinction coefficient or slower chromophore maturation in the cellular environment. Owing to the lack of a mechanistic understanding of the residues involved in chromophore maturation or enhancement in extinction coefficient, we opted for multiple rounds of random (i.e. "error-prone" PCR) mutagenesis to increase the brightness of the C4 mutant.

Error-prone PCR-based selection generated C4PB-12 mutant which differs from C4 by the following mutations: N98K and K166R. Photophysical characterizations of the C4PB-12 mutant showed that it has longer lifetime compared to mCherry (2.22 ns vs. 1.72 ns). However, the dark state conversion and ground state recovery time of this mutant was significantly higher (Table 1, main text). Resultingly, there was significant amount of dark state fraction in this mutant. Therefore, we named C4PB-12 as mCherry-d ("d" stands for dark).

Table S1: Mutations introduced in mCherry-d related to mCherry. The numbering is based on mCherry crystal structure (PDB: 2H5Q).

| residue no | 16 | 17 | 70 | 98 | 99 | 161 | 163 | 166 | 175 | 197 |
|---|---|---|---|---|---|---|---|---|---|---|
| mCherry | V | H | K | N | F | I | Q | K | A | I |
| **C4PB-12 (mCherry-d)** | T | R | R | K | S | M | M | R | W | R |

### S6. Experimental methods for directed evolution of mCherry-d and its photophysical characterizations.

#### Library generation and yeast cell growth

**Template construction:** The original DNA templates for mCherry were amplified using gene-specific primers and cloned into pDonr221 vector using the Gateway recombination system from Life Technologies. The forward primer included a recombination recognition sequence (attB1), a Shine-Dalgarno sequence for prokaryotic expression, a BamH1 restriction endonuclease site, a Kozak sequence for mammalian expression, and a complementary sequence to the FP. The reverse primer included a complementary sequence to the FP, a stop codon, an EcoR1 restriction endonuclease site, and an attB2 recombination recognition sequence. The FP was then cloned into the pYestDest52 vector using the Gateway LR reaction. After confirming the sequences, the FP/pYestDest52 plasmids were used as templates for library construction.

**Site directed libraries:** The QuikChange site-directed mutagenesis kit (Agilent) was used for introducing point mutations or single amino acid switches. PfuTurboDNA polymerase and a thermocycler (BioRad) were employed for the mutagenesis, ensuring high-fidelity replication of both plasmid strands without displacing the mutant oligonucleotide primers. The procedure involved using a supercoiled double-stranded DNA vector containing the target fluorescent protein (FP) along with synthetic oligonucleotide primers carrying the desired mutation. Through temperature cycling, the primers were extended by PfuTurboDNA polymerase, leading to the generation of a mutated plasmid with staggered nicks. DpnI treatment was then applied to digest the parental DNA template and select for mutation-containing synthesized DNA. The nicked vector DNA containing the desired mutations was transformed into E. coli (Top10). For the creation of libraries with multiple site-directed targets, the SOE (Splicing Overlap Extension) reaction was employed. Primers were designed to introduce the desired mutations, and a series of PCR steps were performed to generate overlapping gene segments, which were then used as template DNA for the amplification of full-length products. The resulting full-length product was extracted from a gel, precipitated, and eluted in a small volume of water. Electroporation was conducted using prepared competent yeast cells (*Saccharomyces cerevisiae* BY4741) and the cut pYestDest52 vector. The cell-DNA-vector mixture was subjected to electroporation under specific conditions.

**Error-prone libraries:** EP-PCR libraries were created using the GeneMorph II Random Mutagenesis kit (Agilent Cat No. 200550). The kit protocol was followed with varying amounts of template DNA and cycles depending on the error rate. A typical error-rate is used that incorporates ~5 mutations (at the nucleotide level) per template. 3, 5 T7 and V5 universal primers (both located on pYesDest52 vector) were used for the amplification. After first round of PCR, the gel-extracted PCR product was used for a second round of PCR to create enough DNA for homologous recombination. After PCR purification the library DNA was isopropanol precipitated and eluted in a few µl of water.

**Protein Purification:** Mutants were subsequently transferred to the pBad-His vector for expression in E. coli and purification using Ni-NTA affinity chromatography, and excess imidazole was removed using SnakeSkin dialysis (Thermofisher) bags with buffer exchange (3X over 24h). Purified proteins were consequently diluted in Tris-HCl buffer (pH 7.4) to reach optical densities of approximately 0.1, ensuring that measurements were within the linear range of the instrument response.

**In-vitro characterization and spectroscopy:** Absorption spectra were collected using a Cary 5000 UV−vis near-IR spectrophotometer in the dual-beam mode. The emission and excitation data were collected using a HORIBA Jobin Yvon Fluorolog-3 FL3-222 instrument. Fluorescence lifetime data was collected using a commercial time-correlated single photon counting (TCSPC) system (Fluoro-time 100, PicoQuant). A 560 nm pulsed laser diode head excitation source and a repetition rate of 5 MHz was used. An emission filter centered at 600 nm (60 nm FWHM) was used to filter the excitation and scattered photons. Protocols for measuring fluorescence quantum yield and extinction coefficients are given here.[11]

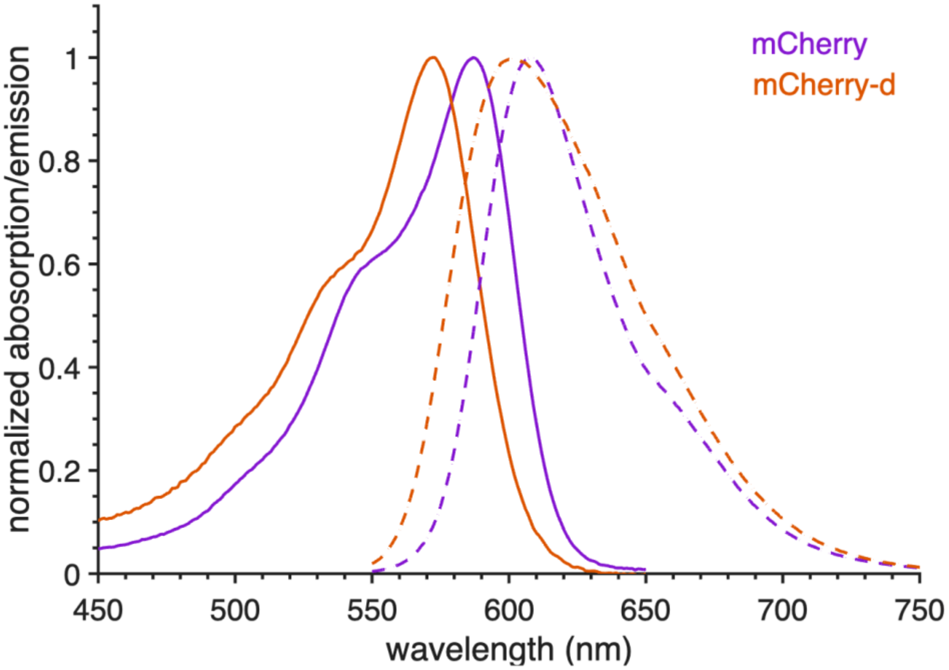


Figure S6.1: Absorption (solid lines) and emission spectra of mCherry (purple) and mCherry-d (orange) variants.

### Microfluidic sorting

**Sorter:** For clone selection, we employed our custom-built single-cell lifetime flow cytometer.[7] The system utilizes a 532-nm laser (Verdi, Coherent, 10W) that is split using multiple beam splitter. The highest power beam passes through an electro-optic modulator (EOM, ThorLabs, EO-AM-NR-C4) to amplitude-modulate the beam at a frequency of 29.5 MHz, we denote this as our lifetime beam. Prior to entering the EOM, the beam is focused with a lens to fit into the EOM aperture. Two polarizers (Newport) and a half-wave plate (Newport) are used to control the power of the lifetime beam after the EOM. The lower power beams and the modulated beam are directed through a 150 mm plano-convex cylindrical lens, which transforms the circular beams into elliptical beams. These elliptical beams, generated by the cylindrical lens, enter the side-port of a commercial inverted microscope (Olympus IX71), pass through a dichroic mirror (Semrock, FF573-Di01–25×36), and are focused into the microfluidic chip (Micronit Technologies, Netherlands) using an air-objective (Olympus, 20x, NA 0.45). The full width at half maximum (FWHM) of the lifetime beam measures ~10 μm and ~60 μm in the minor and major axes of its elliptical spatial mode, respectively. The dimensions of the other beam are similar. Epifluorescence emitted from cells expressing RFPs is separated from the excitation beams using a band-pass filter (Semrock, FF01–629/56–25). These beams are then spatially separated using mirrors and slits and collected by red-wavelength sensitive photomultiplier tubes (PMT, Hamamatsu R9880U-20).The entire setup is controlled using a target and a host computer system operating on LabView 2012 programs.[12]

**Sorting:** The yeast or bacterial cell culture is freshly grown from a stored stock and expressed transiently. A 0.5 ml volume of stored culture media is added to 10 mL solution of growth media (yeast nitrogen base, ammonium sulphate, dextrose) and grown for 8 hrs. for yeast, and for bacteria the 2XYT media with Ampicillin is used. Next, 0.5 ml of this freshly grown cell culture is added to 10 mL solution of induction media (yeast nitrogen base, ammonium sulphate, galactose, raffinose), in case of bacteria 20% arabinose is added to the same. Cells are screened or sorted 17–20 hrs. after induction. During growth and expression, the cultures are incubated at 30°C for yeast and 28°C for bacteria and constantly shaken at 250 rpm. These cells are diluted (10–20 fold) with the blank media (yeast nitrogen base, ammonium sulphate) containing 14% OptiPrep (60% weight/volume iodixanol in water), and subsequently filtered using a 40 µm filter to remove cell debris prior to loading into the microfluidic chip.

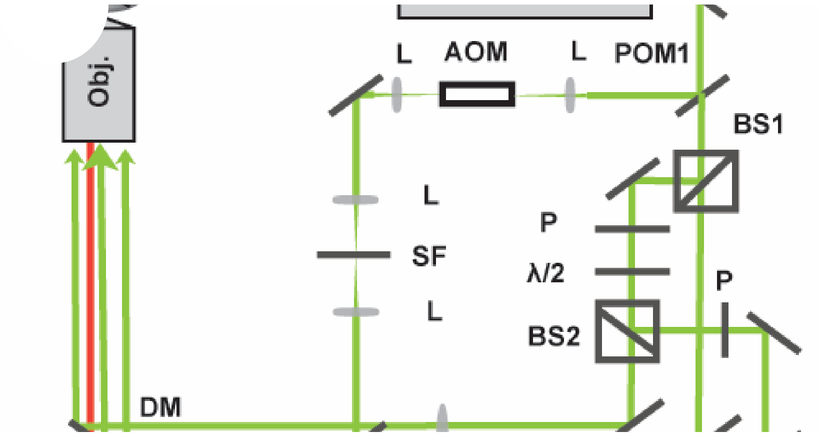


Figure S6.2: The schematic for the microfluidic sorter used in selecting mCherry-d.

### S7. Ground state recovery (GSR) measurement.

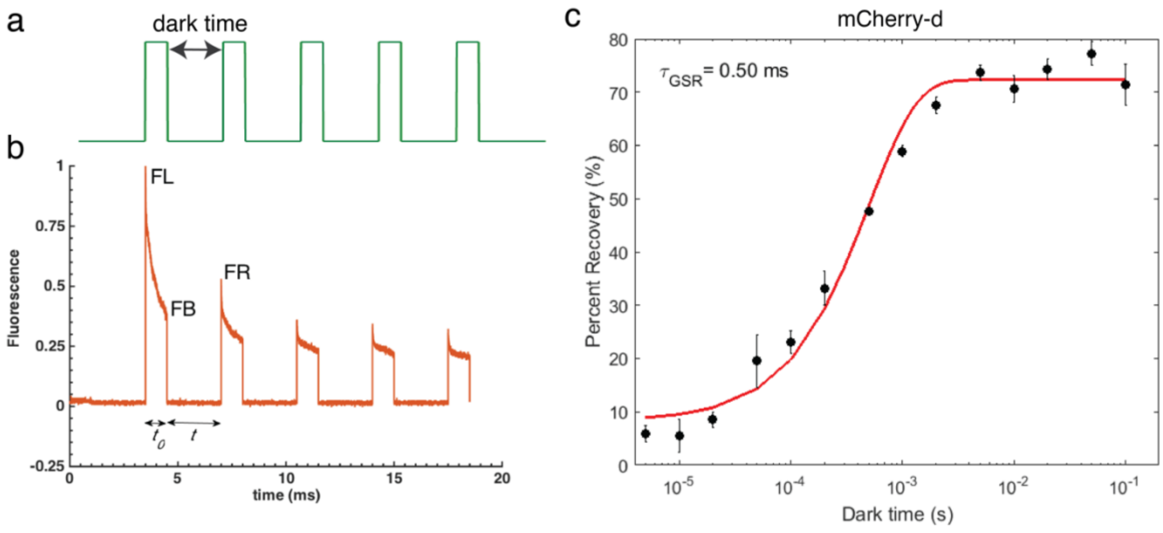


Figure S7: **Ground state recovery measurements of mCherry-d**. (a) Cartoon of laser excitation profile for GSR time measurements with 2 ms of pulsewidth and varying inter-pulse delay or dark time. (b) Typical fluorescence measured from such type of pulsed laser excitation. Fl, FB and FR are used to calculate percent recovery at a particular dark time using Eqn. 25. (c) Percent recovery as a function of different dark time shown in black dots. The red line is the exponential fit. The error-bars in percent recovery times are standard deviation from different immobilized cells.

The details of the ground state recovery measurements can be found here.[13] Briefly, the yeast cells expressing mCherry-d was irradiated with a 532 nm laser pulse sequence similar to shown in Figure S7 a. The exposure time of the pulse were 2 ms and the inter-pulse delays were varied from 5 µs to 100 ms. The percent recovery measured from the fluorescence traces are obtained as:

$$percent\ recovery = \frac{FR-FB}{FL-FB} \times 100 \qquad (25)$$

FR, FB and FL are shown in Figure S7 b. Finally, ground state recovery time-constant ($\tau_{GSR}$) was obtained fitting the percent recovery vs. dark time plot with single exponential function with the form:

$$percent\ recovery = a - b * \exp\left(-t/\tau_{GSR}\right) \qquad (26)$$

where, *a* and *b* are constants. *t* is dark time.

### S8. Measurement of dark state conversion (DSC).

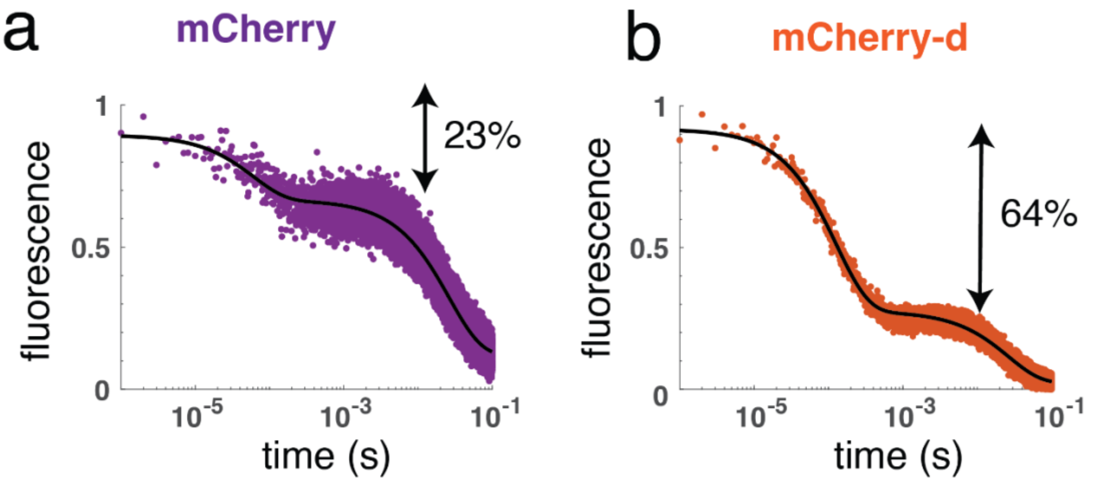


Figure S8: Fluorescence traces obtained from (a) mCherry (purple dots) and (b) mCherry-d (orange dots). The solid black lines are the tri-exponential fits. DSC amplitudes, i.e., the amplitudes of the fast components for mCherry and mCherry-d were 23% and 64% respectively.

For the measurements of DSC time-constants, the fluorescent proteins expressing in yeast cells were irradiated with a 561 nm laser at 5 kW/cm$^2$ in a similar set-up as in.[13] Figure S8, the resulting fluorescence traces from such excitation. The traces contained 3 components: a very fast decay at 1-100 μs timescale, then a flat region which is followed by a slow decay in ms timescale. To obtain the percent DSC, the fluorescence traces were fit to a three-exponential function. Higher weights were assigned to 0-1 ms timescale to accurately capture the DSC process. The fast $\mu s$ time-scale component and its corresponding amplitude from the fits are decay constants ($k_d$) and DSC amplitude ($a_{dsc}$), respectively. Table S3 displays the fitting results of the mean $a_{dsc}$ and $k_d$.

Table S2: DSC amplitude and decay constants for mCherry and mCherry-d. n = biological replicates. Values are average of the replicates, and the errors are standard deviation.

| | $a_{dsc}$ (%) | $k_d$ (μs) |
|---|---|---|
| mCherry | 23 ± 1.2 (n=5) | 59 ±16 (n=5) |
| mCherry-d | 64 ± 3.3 (n=4) | 138 ±18 (n=4) |

Finally, Eqn. 20 was used to quantify the DSC time-constants with known values of $k_{gsr}$ and $q$ obtained from independent measurements.

### S9. Analysis of photobleaching decays in mCherry and mCherry-d.

For the photobleaching experiments, yeast cells expressing RFPs were illuminated for 2 s using a 532-nm laser at 1, 5, and 10 kW/cm$^2$ intensities. For each sample and intensity, three independent measurements were taken. The initial sub-ms fluorescence decay largely due to dark state conversion and therefore discarded for photobleaching analyses and fits. The remaining background-corrected fluorescence decay traces were fit to a bi-exponential function of the form of Eqn. 5 (main text). For each sample, decays with different intensities are globally fit with a bi-exponential function where $\tau_1$ is kept as a shared variable. On the other hand, the amplitudes ($a_1$, $a_2$) and $\tau_2$ are kept as intensity data specific.

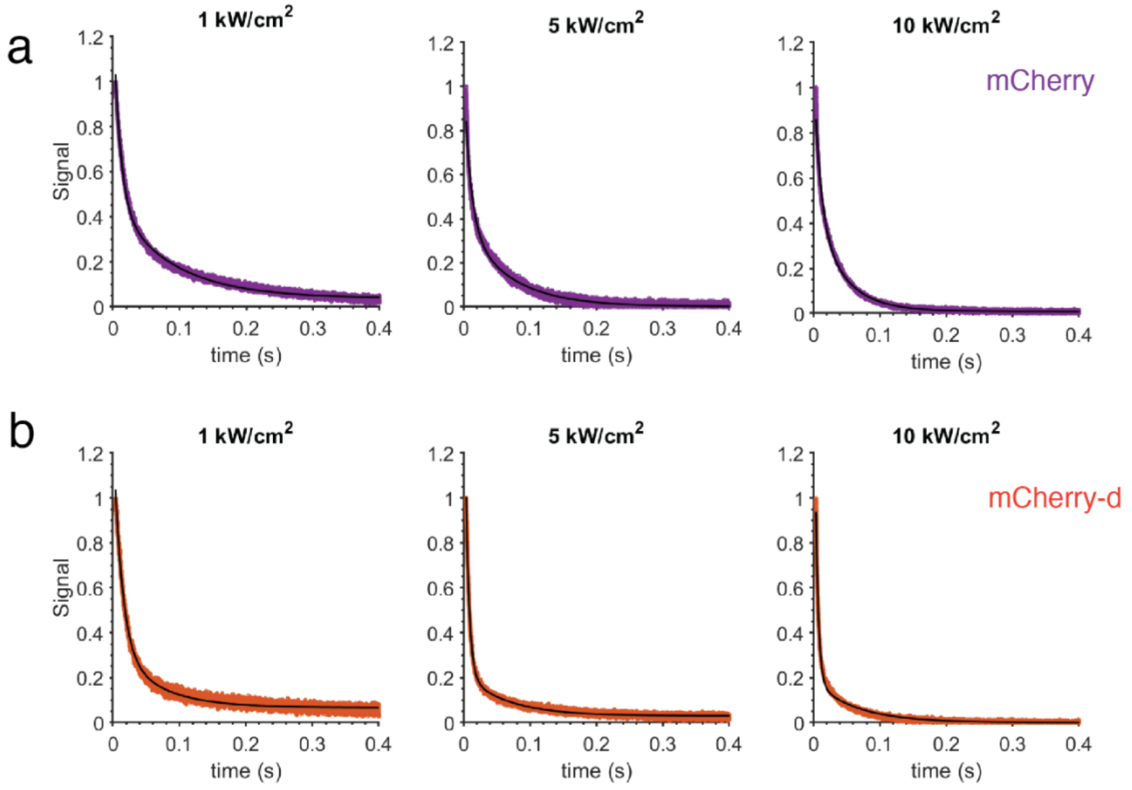


Figure S9.1: **Fits of the continuous photobleaching decays.** Photobleaching traces of (a) mCherry (purple) and (b) mCherry-d (orange) at different laser intensities under continuous illumination mode. The bi-exponential fits are shown in the black lines. The initial 4 ms of traces those largely contained dark state conversion are discarded for photobleaching analysis. The fit values are shown in Table 2, main text.

### S10. Molecular dynamics simulations of mCherry and mCherry-d

The crystal structure of mCherry was obtained from the Protein Data Bank (PDB ID: 2H5Q).[14] To create the starting structure for mCherry-d the crystal structure was modified at each mutated position with Chimera ver 1.19, with rotamers chosen from the Dunbrack rotamer database.[15,16] These structures were checked and protonated with MolProbity.[17,18] Protonation states were tested at a pH of 7.0 using propka3, though all residues maintained their standard protonation states according to their pKa.[19]

Each system was neutralized to a net charge of zero with Na+ ions (4 in mutant, 7 in WT), and solvated with TIP3P water such that there was a minimum of 15 Å from the surface of the protein to the side of the box.[20] The ff14SB force field was used for the protein structure, and parameters for the chromophore in its phenolate form with the deprotonated acylimine nitrogen typical of RFPs were generated with AutoParams as in previous work.[21–23] Each system was subjected to an initial minimization, density equilibration at 10 K, temperature equilibration to 300 K, and a final density equilibration before production MD.

Production MD was run in Amber 20 in the NVT ensemble. SHAKE and RATTLE were used to constrain the hydrogens and allow for a 2 fs timestep.[24,25] The GPU-accelerated pmemd.cuda module as implemented in Amber 20 was used with a 10 Å cutoff for the nonbonded interactions and smooth particle mesh Ewald for the electrostatics.[26,27] Each system was run for 500 ns of production in duplicate.

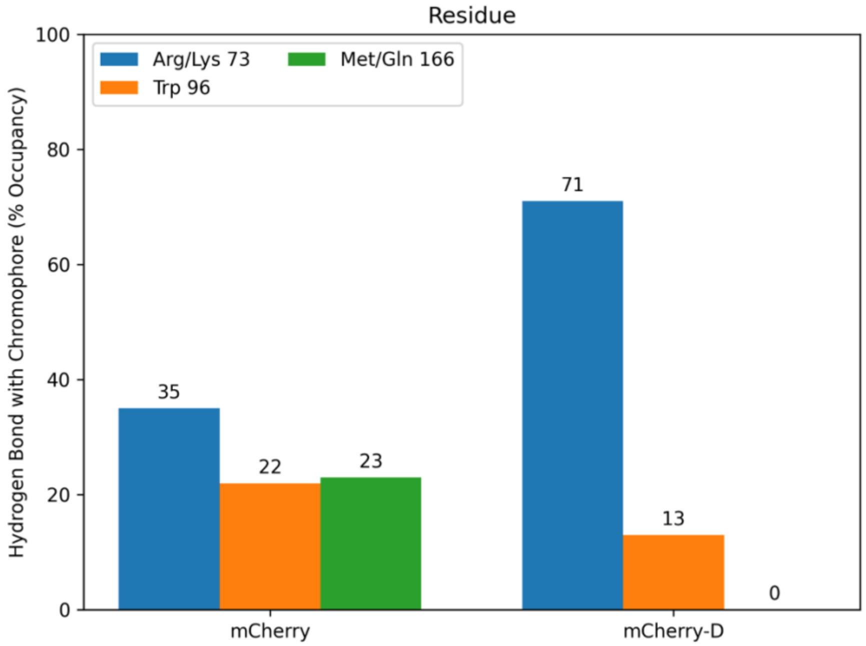


Figure S10: Percent occupancy of hydrogen bond with chromophore for select amino acid residues in mCherry and mCherry-d. The numbering of residues in this plot and Table below are different. The corresponding residues in the main texts (based on PDB 2H5Q ) are: 73 → 70, 166→163 and 96→93.

**Hydrogen Bonding Table**

**Notes:** The hydrogen bonding occupancies are summed over consistent interactions. So, in the case of e.g. arginine residues, there are ~6 possible hydrogen bonds. 100% means there is at least 1 hydrogen bond 100% of the time, 200% would mean 2 hydrogen bonds 100% of the time, etc.

The differences are relative to mCherry-d. A positive percentage means an increase in hydrogen bond presence in mCherry-d, and a negative percentage means a decrease.

| Acceptor Residue | Donor Residue | mCherry % Occupancy | mCherry-d % Occupancy | Difference (mCherry-d – mCherry) |
|---|---|---|---|---|
| 1 | 191 | 0.53% | 25.46% | 24.93% |
| 1 | 233 | 0.00% | 19.86% | 19.86% |
| 1 | 234 | 0.00% | 14.22% | 14.22% |
| 2 | 234 | 0.00% | 14.51% | 14.51% |
| 3 | 6 | 2.24% | 24.32% | 22.08% |
| 3 | 92 | 18.93% | 0.00% | -18.93% |
| 4 | 7 | 28.25% | 45.46% | 17.21% |
| 5 | 7 | 12.67% | 0.45% | -12.22% |
| 6 | 6 | 18.27% | 3.15% | -15.12% |
| 7 | 7 | 20.95% | 1.46% | -19.49% |
| 7 | 9 | 14.80% | 4.16% | -10.64% |
| 7 | 84 | 0.00% | 8.82% | 8.82% |
| 10 | 13 | 39.95% | 48.00% | 8.05% |
| 21 | 70 | 0.00% | 25.78% | 25.78% |
| 22 | 24 | 0.20% | 65.74% | 65.54% |
| 22 | 33 | 65.90% | 94.07% | 28.17% |
| 22 | 35 | 0.00% | 61.33% | 61.33% |
| 39 | 48 | 161.98% | 148.13% | -13.85% |
| 41 | 46 | 88.28% | 72.51% | -15.77% |
| 47 | 72 | 111.02% | 42.55% | -68.47% |
| 65 | 69 | 23.72% | 59.88% | 36.16% |
| 66 | 69 | 47.47% | 25.90% | -21.57% |
| 70 | 72 | 6.33% | 55.88% | 49.55% |
| **71 (CRO)** | **73 (70 in 2H5Q)** | **34.95%** | **71.32%** | **36.37%** |
| **71(CRO)** | **96 (93 in 2H5Q)** | **21.72%** | **12.89%** | **-8.83%** |
| 74 | 87 | 16.36% | 28.61% | 12.25% |
| 76 | 87 | 25.75% | 15.48% | -10.27% |
| 78 | 84 | 52.79% | 61.21% | 8.42% |
| 85 | 89 | 49.77% | 16.38% | -33.39% |
| 86 | 89 | 59.36% | 88.15% | 28.79% |

| | | | | |
|---|---|---|---|---|
| 92 | 92 | 47.88% | 57.44% | 9.56% |
| 101 | 179 | 124.59% | 115.18% | -9.41% |
| 102 | 104 | 0.10% | 97.19% | 97.09% |
| 102 | 106 | 51.10% | 67.77% | 16.67% |
| 103 | 177 | 80.66% | 104.92% | 24.26% |
| 110 | 98 | 144.17% | 132.56% | -11.61% |
| 111 | 96 | 34.90% | 52.03% | 17.13% |
| 111 | 97 | 20.89% | 12.36% | -8.53% |
| 112 | 96 | 93.44% | 113.65% | 20.21% |
| 114 | 94 | 128.64% | 118.27% | -10.37% |
| 114 | 187 | 64.12% | 51.77% | -12.35% |
| 115 | 122 | 109.83% | 95.79% | -14.04% |
| 117 | 120 | 82.91% | 72.23% | -10.68% |
| 123 | 21 | 0.00% | 42.54% | 42.54% |
| 123 | 22 | 20.38% | 6.75% | -13.63% |
| 123 | 72 | 38.45% | 0.22% | -38.23% |
| 126 | 24 | 34.41% | 43.95% | 9.54% |
| 127 | 128 | 28.83% | 13.14% | -15.69% |
| 128 | 24 | 54.16% | 23.99% | -30.17% |
| 128 | 26 | 5.60% | 18.52% | 12.92% |
| 128 | 31 | 13.44% | 32.44% | 19.00% |
| 130 | 131 | 40.83% | 21.76% | -19.07% |
| 144 | 169 | 79.25% | 130.27% | 51.02% |
| 146 | 169 | 2.51% | 17.82% | 15.31% |
| 146 | 201 | 12.40% | 0.53% | -11.87% |
| 147 | 166 | 26.57% | 0.00% | -26.57% |
| 147 | 167 | 47.98% | 56.52% | 8.54% |
| 147 | 169 | 13.60% | 89.99% | 76.39% |
| 147 | 175 | 77.89% | 54.13% | -23.76% |
| 149 | 165 | 87.36% | 2.30% | -85.06% |
| 149 | 200 | 48.63% | 91.86% | 43.23% |
| 150 | 197 | 38.16% | 6.27% | -31.89% |
| 150 | 231 | 21.32% | 0.00% | -21.32% |
| 151 | 73 | 35.37% | 191.07% | 155.70% |
| 151 | 184 | 82.27% | 25.25% | -57.02% |
| 151 | 198 | 152.10% | 129.34% | -22.76% |
| 151 | 200 | 0.00% | 111.74% | 111.74% |
| 152 | 163 | 209.52% | 91.78% | -117.74% |
| 152 | 197 | 0.71% | 19.55% | 18.84% |
| 152 | 230 | 0.00% | 21.87% | 21.87% |
| 154 | 161 | 142.40% | 154.94% | 12.54% |
| 154 | 163 | 35.63% | 43.76% | 8.13% |
| 156 | 157 | 8.76% | 17.55% | 8.79% |

| | | | | |
|---|---|---|---|---|
| 162 | 182 | 134.03% | 114.78% | -19.25% |
| **166 (163 in 2H5Q)** | **71 (CRO)** | **23.12%** | **0.00%** | **-23.12%** |
| 166 | 167 | 21.37% | 0.00% | -21.37% |
| 167 | 177 | 138.92% | 173.25% | 34.33% |
| 167 | 234 | 14.21% | 0.00% | -14.21% |
| 181 | 99 | 92.86% | 83.36% | -9.50% |
| 181 | 182 | 0.01% | 16.55% | 16.54% |
| 184 | 73 | 0.12% | 30.25% | 30.13% |
| 185 | 95 | 149.52% | 159.98% | 10.46% |
| 187 | 92 | 6.54% | 18.62% | 12.08% |
| 187 | 93 | 58.11% | 9.29% | -48.82% |
| 188 | 7 | 9.80% | 0.00% | -9.80% |
| 188 | 8 | 23.96% | 14.01% | -9.95% |
| 188 | 88 | 12.94% | 3.81% | -9.13% |
| 191 | 3 | 0.31% | 70.82% | 70.51% |
| 191 | 192 | 5.38% | 49.98% | 44.60% |
| 196 | 78 | 41.65% | 60.97% | 19.32% |
| 197 | 223 | 235.97% | 95.25% | -140.72% |
| 197 | 226 | 0.00% | 10.18% | 10.18% |
| 199 | 221 | 114.40% | 136.00% | 21.60% |
| 199 | 223 | 4.26% | 18.47% | 14.21% |
| 201 | 203 | 43.86% | 34.74% | -9.12% |
| 201 | 219 | 62.40% | 81.00% | 18.60% |
| 203 | 204 | 2.02% | 10.49% | 8.47% |
| 203 | 217 | 141.40% | 124.70% | -16.70% |
| 203 | 219 | 140.68% | 5.54% | -135.14% |
| 216 | 48 | 51.75% | 62.25% | 10.50% |
| 216 | 49 | 118.76% | 107.59% | -11.17% |
| 218 | 46 | 38.58% | 67.39% | 28.81% |
| 218 | 47 | 198.23% | 187.08% | -11.15% |
| 218 | 73 | 34.99% | 52.40% | 17.41% |
| 219 | 44 | 6.03% | 39.62% | 33.59% |
| 220 | 45 | 46.62% | 67.46% | 20.84% |
| 221 | 223 | 0.89% | 57.10% | 56.21% |
| 223 | 230 | 63.05% | 0.00% | -63.05% |
| 223 | 231 | 74.62% | 0.04% | -74.58% |
| 224 | 81 | 2.60% | 21.79% | 19.19% |
| 224 | 226 | 14.14% | 5.49% | -8.65% |
| 224 | 227 | 35.70% | 0.97% | -34.73% |
| 224 | 228 | 12.24% | 0.00% | -12.24% |
| 224 | 230 | 8.69% | 0.15% | -8.54% |
| 225 | 227 | 0.08% | 12.82% | 12.74% |

| | | | | |
|---|---|---|---|---|
| 226 | 229 | 14.14% | 0.42% | -13.72% |
| 226 | 230 | 44.39% | 1.75% | -42.64% |
| 230 | 234 | 14.84% | 42.26% | 27.42% |
| 233 | 6 | 0.00% | 17.09% | 17.09% |

(11) Manna, P.; Hung, S.-T.; Mukherjee, S.; Friis, P.; Simpson, D. M.; Lo, M. N.; Palmer, A. E.; Jimenez, R. Directed Evolution of Excited State Lifetime and

Brightness in FusionRed Using a Microfluidic Sorter. *Integr. Biol.* **2018**, *10* (9), 516–526.
(12) Mukherjee, S. Spectroscopic Evaluation of Excited State Depopulation in Red Fluorescent Proteins Developed Using Fluorescence Lifetime Selections. PhD Thesis, University of Colorado at Boulder, 2022.
(13) Manna, P.; Jimenez, R. Time and Frequency-Domain Measurement of Ground-State Recovery Times in Red Fluorescent Proteins. *J. Phys. Chem. B* **2015**, *119* (15), 4944–4954.
(14) Shu, X.; Shaner, N. C.; Yarbrough, C. A.; Tsien, R. Y.; Remington, S. J. Novel Chromophores and Buried Charges Control Color in mFruits,. *Biochemistry* **2006**, *45* (32), 9639–9647. https://doi.org/10.1021/bi060773l.
(15) Pettersen, E. F.; Goddard, T. D.; Huang, C. C.; Couch, G. S.; Greenblatt, D. M.; Meng, E. C.; Ferrin, T. E. UCSF Chimera: A Visualization System for Exploratory Research and Analysis. *J Comput Chem* **2004**, *25* (13), 1605–1612. https://doi.org/10.1002/jcc.20084.
(16) Dunbrack, R. L.; Karplus, M. Backbone-Dependent Rotamer Library for Proteins. Application to Side-Chain Prediction. *J. Mol. Biol.* **1993**, *230* (2), 543–574. https://doi.org/10.1006/jmbi.1993.1170.
(17) Chen, V. B.; Arendall, W. B.; Headd, J. J.; Keedy, D. A.; Immormino, R. M.; Kapral, G. J.; Murray, L. W.; Richardson, J. S.; Richardson, D. C. *MolProbity* : All-Atom Structure Validation for Macromolecular Crystallography. *Acta Crystallogr. D Biol. Crystallogr.* **2010**, *66* (1), 12–21. https://doi.org/10.1107/S0907444909042073.
(18) Williams, C. J.; Headd, J. J.; Moriarty, N. W.; Prisant, M. G.; Videau, L. L.; Deis, L. N.; Verma, V.; Keedy, D. A.; Hintze, B. J.; Chen, V. B.; Jain, S.; Lewis, S. M.; Arendall, W. B.; Snoeyink, J.; Adams, P. D.; Lovell, S. C.; Richardson, J. S.; Richardson, D. C. MolProbity: More and Better Reference Data for Improved All-Atom Structure Validation. *Protein Sci. Publ. Protein Soc.* **2018**, *27* (1), 293–315. https://doi.org/10.1002/pro.3330.
(19) Olsson, M. H. M.; Søndergaard, C. R.; Rostkowski, M.; Jensen, J. H. PROPKA3: Consistent Treatment of Internal and Surface Residues in Empirical pKa Predictions. *J. Chem. Theory Comput.* **2011**, *7* (2), 525–537. https://doi.org/10.1021/ct100578z.
(20) Jorgensen, W. L.; Chandrasekhar, J.; Madura, J. D.; Impey, R. W.; Klein, M. L. Comparison of Simple Potential Functions for Simulating Liquid Water. *J. Chem. Phys.* **1983**, *79* (2), 926–935. https://doi.org/10.1063/1.445869.
(21) Maier, J. A.; Martinez, C.; Kasavajhala, K.; Wickstrom, L.; Hauser, K. E.; Simmerling, C. ff14SB: Improving the Accuracy of Protein Side Chain and Backbone Parameters from ff99SB. *J. Chem. Theory Comput.* **2015**, *11* (8), 3696–3713. https://doi.org/10.1021/acs.jctc.5b00255.
(22) Hix, M. A.; Walker, A. R. AutoParams: An Automated Web-Based Tool To Generate Force Field Parameters for Molecular Dynamics Simulations. *J. Chem. Inf. Model.* **2023**, *63* (20), 6293–6301. https://doi.org/10.1021/acs.jcim.3c01049.
(23) Murphy, A. R.; Hix, M. A.; Walker, A. R. Exploring the Effects of Mutagenesis on FusionRed by Using Excited-State QM/MM Dynamics and Classical Force Field Simulations. *ChemBioChem* **2023**, *24* (12), e202200799. https://doi.org/10.1002/cbic.202200799.

(24) Ryckaert, J.-P.; Ciccotti, G.; Berendsen, H. J. C. Numerical Integration of the Cartesian Equations of Motion of a System with Constraints: Molecular Dynamics of n-Alkanes. *J. Comput. Phys.* **1977**, *23*, 327–341. https://doi.org/10.1016/0021-9991(77)90098-5.
(25) Miyamoto, S.; Kollman, P. A. Settle: An Analytical Version of the SHAKE and RATTLE Algorithm for Rigid Water Models. *J. Comput. Chem.* **1992**, *13* (8), 952–962. https://doi.org/10.1002/jcc.540130805.
(26) Salomon-Ferrer, R.; Götz, A. W.; Poole, D.; Le Grand, S.; Walker, R. C. Routine Microsecond Molecular Dynamics Simulations with AMBER on GPUs. 2. Explicit Solvent Particle Mesh Ewald. *J. Chem. Theory Comput.* **2013**, *9* (9), 3878–3888. https://doi.org/10.1021/ct400314y.
(27) Darden, T.; York, D.; Pedersen, L. Particle Mesh Ewald: An N·log(N) Method for Ewald Sums in Large Systems. *J. Chem. Phys.* **1993**, *98* (12), 10089–10092. https://doi.org/10.1063/1.464397.
